# Gaze dynamics prior to navigation support hierarchical planning

**DOI:** 10.1101/2025.01.16.633460

**Authors:** Jeremy Gordon, John Chuang, Giovanni Pezzulo

## Abstract

The task of planning future actions in the context of an uncertain world results in massive state spaces that preclude exhaustive search and other strategies explored in the domains of both human decision-making and computational agents. One plausible solution to this dimensionality explosion is to decompose the task into subgoals that match the information geometry of the task at hand. However, how individuals identify a productive hierarchy, and perceive and select subgoals suitable to planning, is not well understood. To investigate this topic, we designed a virtual-reality based behavioral experiment which collected eye movements during a pre-navigation planning phase. By capturing gaze dynamics correlated with the simulative processes used in planning, we were able to identify the spatiotemporal evolution of visual search under uncertainty. Our results highlight gaze dynamics indicative of a search process that exhibits hierarchical structure. These include a decreasing trend seen in gaze distance from origin and a broad to narrow shift (with reducing saccade distances and longer fixation durations) as plans are established. In line with prior work, critical tiles to which landscape connectivity is most sensitive were the strongest predictors of visual attention. We also find that deeper planning was correlated with success only on the most complex maps (e.g. those with a larger number of information-nodes, higher branching factor, and more forks, according to an info-graphical map analysis). This study highlights the role of embodied visual search during planning, and the skill-dependence of the specific subgoals and hierarchical decomposition used which unlocked successful performance.

## Introduction

When planning future actions, e.g. finding the best route to a novel destination or the best sequencing for our daily tasks, we are often confronted with very large state spaces that we cannot search exhaustively. These exist among a class of problems we frequently encounter as humans which involves searching for a sequence of candidate actions likely to reliably result in a desired “goal” state, while also contending with limited (cognitive and motor) resources. Prospective search problems of this type are common in the realm of spatial navigation, but also apply to much more abstract domains of cognition [1–3].

From a formal perspective, searching in large state spaces is different from searching in small ones that permit exhaustive analysis, and as a result, requires a different class of algorithms [4]. Indeed, in discrete state spaces with known dynamics, it is possible to use simple decision trees that capture a complete set of possible action sequences, to identify paths resulting in a desired state, as well as an evaluation of their relative efficiency (e.g. by weighted length). However, in large state spaces (including tasks where states are continuous rather than discrete), the combinatorial nature of possible action sequences results in a dimensionality explosion that makes exhaustive search intractable. Even when loops are excluded (to avoid an infinite search space), the number of sequences to check exceeds the computational capabilities of both biological and artificial problem solvers. In machine learning, the problem of searching in large state spaces has lead to the development of various effective techniques to *sample* the most promising candidate actions [4]. For example, Monte Carlo Tree Search (MCTS) algorithms, when coupled with deep neural networks, provide effective solutions to challenging games such as chess and Go [5]. In parallel, the identification of algorithms and heuristics organisms use to overcome these challenges has become a rich target of inquiry for cognitive science and psychology [6, 7].

When faced with large search problems, living organisms (as well as artificial systems) must contend with the *breadth-depth dilemma*, in which the problem solver must decide how to direct limited resources (e.g. exploratory actions which consume time and energy). Typically, spending more time reasoning about (simulating) sequences within a particular region of a state space offers more precise estimates of its value (i.e. in pathfinding, the likelihood of particular positions being part of an efficient route to destination), at the cost of reduced precision in other regions. A number of studies have attempted to model and empirically measure human handling of this dilemma. In a study in which participants had to select between 121 options that provided different rewards, Wu et al. found that people sometimes adopt a sampling strategy similar to those used in machine learning and that they are able to use spatial structure (when present) to generalize effectively [8]. In a problem solving study resembling the traveling salesman problem, Eluchans et al. found that people adapt their planning strategy to the minimal depth required by the problem [9]. In a patch foraging-like experiment where the allocation of samples needed to be determined prior to sample feedback, Moreno et al. found that individuals prefer breadth when capacity is low, and depth when capacity is high [10]. Interestingly, even when sufficient capacity is available to sample from all states, optimal behavior involved ignoring the majority of states in order to more deeply explore a subset. In a related study, the number of states sampled increased with capacity approximately according to a power law (with exponent 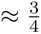) [11].

Other studies have shown that when planning routes in novel mazes, people focus their attention and search resources on *salient* locations or landmarks, which provide more information gain. For example, during virtual navigation, people display more deliberative behavior (as indexed by a behavioral index called *vicarious trial-and-error* (VTE) and initially studied by Tolman during rodent navigation [12]) in locations of the maze where making optimal decisions required using more information [13, 14]. When people have access to the search space (e.g., a map of the maze) prior to navigation and can therefore form a prospective plan, they again focus their resources for encoding with high fidelity only the subset of map elements that are relevant for their (optimal) plan, therefore constructing a simplified mental representation of the problem [15]. Similarly, another study suggests that people adaptively *compress* simulations of potential routes during prospective route planning [16], using mechanisms that are potentially analogous to the (time-compressed) ‘(p)replay’ of experience found in the rodent hippocampus that might support spatial planning [17–24].

Another useful approach to search in large state spaces consists of decomposing the problem into smaller and more manageable sub-problems, thus using a hierarchical approach to planning as opposed to “flat” tree search. Various studies indicate that people use such hierarchical planning strategies when the environment permits hierarchical decomposition [25–29]. During navigation in a maze structured similarly to the “Hanoi tower” problem [30], participants faced with a choice between two paths of the same length preferred the path that crossed fewer community boundaries (as identified by graph theoretic measures), which is “shorter” as quantified by information-theoretic measures that reflect the hierarchical clustering of space [26, 31]. Another study reported that people spontaneously organize space into clusters that support hierarchical planning and subgoaling [32]. A neuroimaging study suggested that when planning routes in a virtual subway network, people exploit the problem structure—and in particular the division into lines and stations—to form hierarchically organized plans [25]. Interestingly, besides hierarchical problem decomposition, individuals often exhibit sensitivity to the structure of the environment in their planning and navigation strategies. For example, a large scale virtual navigation study showed that people are better at wayfinding in mazes having the same general structure as the place where they grew up [33].

Many of the studies reviewed above measure navigation behavior to infer planning strategies. However, planning is a largely covert process, implying that key dynamics may be missed by studying only wayfinding decisions. An alternative approach to study planning in large state spaces consists of recording eye movements while people visually scan the environment prior to beginning navigation. Given its flexibility and speed, the visual system affords a rapid virtual simulation of candidate routes prior to actual movement. Therefore, looking at eye movements may provide privileged access to the covert stages of planning [34, 35]. Along these lines, in a recent study of visual search prior to a navigation task, Zhu et al. showed that participants’ eyes made fast ‘sweeps’ that traced the trajectories that participants later selected, and that gaze targets balanced fixations on goal locations and critical transitions (obstacles or openings) in the environment—findings which resemble results in neural replay [36]. Other studies of eye movements during movement preparation showed that visual attention was selectively allocated to goal-related information, such as the location of movement targets [37] and the planned trajectory around obstacles [38]. Another study of visual search in humans has shown that eye movements themselves exhibit behavior consistent with multi-step planning (in this case, planning of a gaze sequence matched to the shape of a given search space), rather than greedy selection of informationally rich or otherwise salient locations [39]. Multi-step planning of eye movements has been reported also during tasks requiring participants to fixate sequences of stationary targets [40] and to find paths in a grid that collect multiple gems without passing through the same node twice [41]. In a primate study, Lakshminarasimhan et al. showed that eye movements could be used to infer latent beliefs such as the location of a hidden goal during virtual spatial navigation [42]. Controlling fixations also showed detrimental effects on performance, indicating the involvement of free visual search in effective navigation.

Taken together, this literature indicates that when planning in large state spaces that cannot be searched exhaustively, people (and other mammals) use a range of effective strategies: they allocate limited resources by privileging information-rich states, they attend to the most salient aspects of the problem, and they split problems into more manageable sub-problems. These and other strategies can be interpreted in terms of a rational use of limited (bounded) resources [43–45, 45–48]. However, despite these achievements, we still lack a comprehensive account of the planning strategies that people adopt when confronted with the exploration-exploitation dilemma inherent to visual exploration of large, novel and uncertain environments. Interestingly, while behavioral studies have found signatures of hierarchical planning during navigation, we still do not know how these hierarchical navigation plans might be established during visual exploration prior to movement.

In this work, we address three main questions. First, we ask whether gaze dynamics prior to movement show signatures of hierarchical planning and subgoaling – perhaps as a way to establish a hierarchical navigation plan. If planning resembles a flat search in a decision tree, as in MCTS, we should see a series of broader-to-finer-grained series of forward gazes, from the start to the potential goal locations. If instead planning is hierarchical, we should find that people first gaze and resolve uncertainty at the highest level (e.g., about goal and subgoal locations) and subsequently focus at the lower level, to select specific routes from one subgoal to the next – or backward, from the final goal to the intermediate subgoals to the start location. Second, in case of hierarchical planning, we ask how people select subgoal locations – and whether subgoal selection reflects critical transition points and information-theoretic principles of optimal plan decomposition, as suggested by theoretical models [26, 30, 31, 49]. Third, we ask which of these behavioral patterns and planning dynamics led to success planning under uncertainty. That is, we aim to answer the question: what sets top-performing planners apart?

To investigate these questions, we designed a novel Virtual Reality-based spatial navigation task where planning is explicitly required, and collected eye tracking data while participants searched the landscape prior to navigating through it. In contrast to standard wayfinding experiments in which only one goal location is present, the maps were designed with multiple candidate goal locations (“caches”), and participants did not know in advance which ones would be filled in with actual goals. This uncertainty is intended to increase planning demands, and align our task with the inherent uncertainty found within most real world planning contexts.

## Methods

### Participants

Participants were undergraduates, graduate students and staff at a university in the United States. The recruitment period for this study was between Nov 12, 2021 and Feb 9, 2022. Participants were invited by email to complete a screening questionnaire. We excluded participants who were deemed high risk for adverse effects of using VR. This includes those who have used VR in the past and experienced severe dizziness or nausea during usage, as well as those with epilepsy, a history of seizures, heart ailments, or those who respond that they are prone to motion sickness. Participants who are left handed or ambidextrous were also excluded in order to ensure standardized analysis of controller motion in the dominant hand. Participants were also required to be 18 years old or older, and speak English (native English was not a requirement).

After giving written informed consent, 40 participants began the study, but 6 had to stop early due to motion sickness. In total, 34 participants completed the study, and form the sample for this analysis. Mean participant age was 21.3 *±* 3.0. 75% were female, 21% were male, and 4% declined to state. 62% identified as Asian, 21% as White, 3% as Black or African American, and 8% identified as multiple races or Other. No minors participated in the study.

### Experimental design

The study used a single-condition within subjects design. Maps were procedurally generated (see Supporting information for details), and balanced between high uncertainty (2 true goals among 6 possible goal locations), and low uncertainty (2 true goals among 3 possible goal locations). The same group of 33 maps was shown to all participants. The order of the first 30 (the procedurally generated set) was randomized uniformly. The final 3 hand-designed maps were shown at the end of the study, in the same order for all participants. See Fig 1 for an overview of study design.

**Figure 1:**
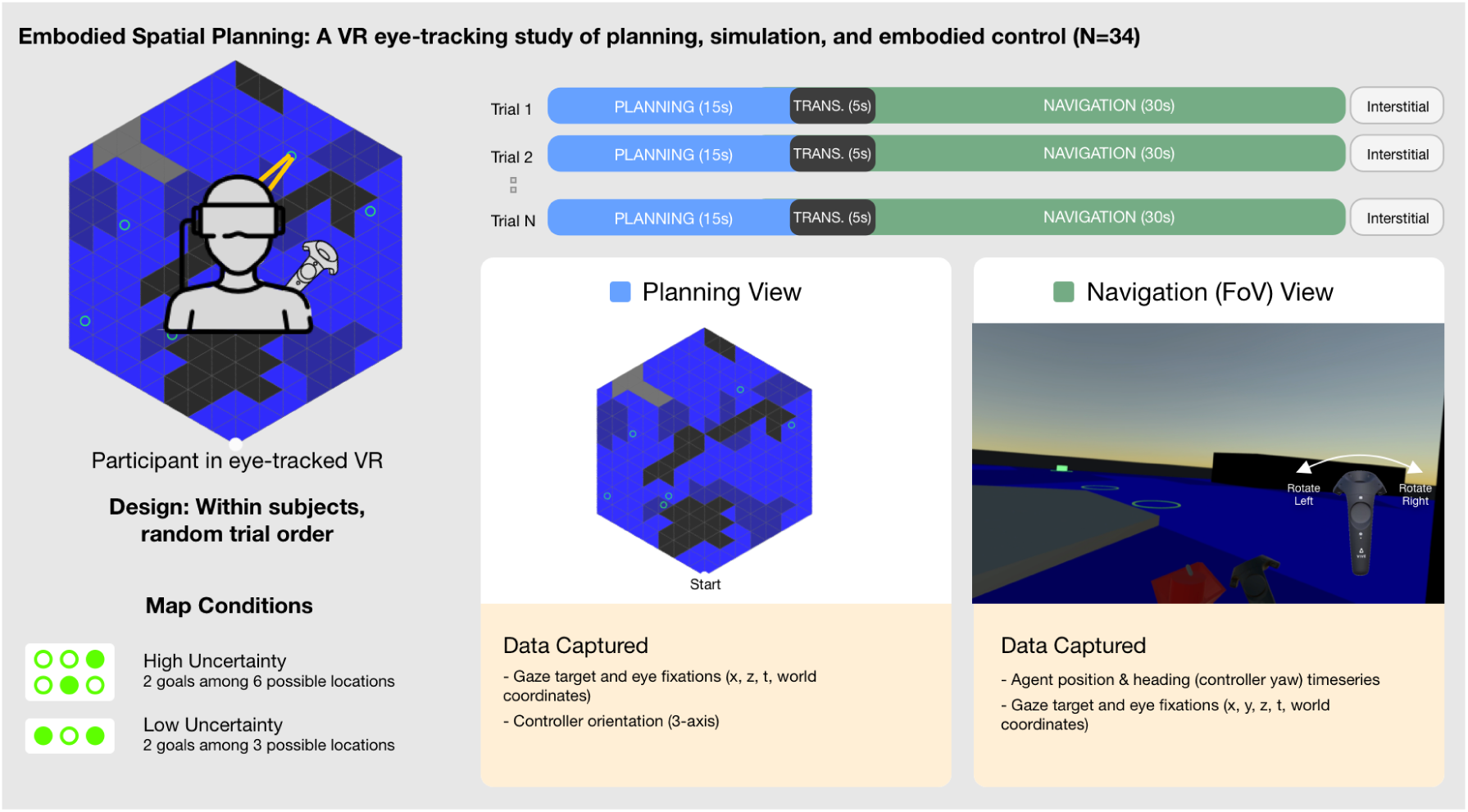
Schematic of experiment design. Participants moved through a series of trials, each split into a planning phase (15s), in which they viewed the map top down, and a navigation phase (30s), in which they moved through the environment with the goal of finding each of two goals. A 5-second transition phase marked the switch between the two main phases. Raw and target-aware eye-tracking data was captured during all three phases. Traces of agent location and orientation were captured during the navigation phase. Each trial presented a unique procedurally generated map in one of two conditions (high or low uncertainty, implemented by the number of goal cache locations). Tile type color key - blue: water, dark blue: muddy water, gray: land, black: obstacle.

### Apparatus

An HTC Vive Pro Eye head mounted display (HMD), which was preconfigured with two Tobii eye-tracking cameras, was the primary apparatus supporting this study. The HMD was driven by an Alienware laptop running experiment scenes in Unity. A single HTC Vive controller was given to each participant to hold in the dominant hand. A “room-scale” setup with two lighthouses positioned on either side of the participant’s seat was used.

### Task & procedure

Prior to beginning the experiment, participants were briefed, signed an informed consent form, and were fitted with the VR headset. An eye-tracking calibration exercise was performed prior to starting the experiment. Then, a series of 3 practice trials began. In each, a simple map was shown to the participant, to ensure familiarity with the interaction and the task. We explained the interaction, meaning of colored tiles, and task objective (to find and collect both goals before time expires) as the participant completed the three practice trials, answering any questions that were raised.

Following the three practice trials, the main experiment begins. Each trial is constituted by a unique map, and progresses in three phases. In the *planning phase* (15 seconds), participants see a top-down view of the landscape in front of them. This map view shows the starting location of a boat—an agent which the participant controls—at the bottom of the map, as well as the colored features of the landscape. Possible goal locations are shown as green rings. Tall obstacles are shown as black tiles, land (which can be seen over but not passed through) is shown as gray tiles, and blue and dark blue tiles indicate water. During the planning phase, the participant is able to view the map, but cannot yet take any navigation action. 3 chimes indicate the approach of the end of the planning phase. Next, in the *transition phase* (5 seconds), the map is removed and participants wait for the beginning of the next phase. In the third and final phase, the *navigation phase* (30 seconds), participants’ view shifts and they are now situated in the landscape just behind the boat they control. They can look in any direction to explore the landscape from the agent’s perspective. By rotating the controller around the vertical axis (yaw), the participant controls the direction of the boat. By holding down the trigger, the participant is able to move the boat forward in its present orientation. In this way, participants are able to navigate the landscape with the aim of reaching and collecting both goals in the map. If the boat contacts either type of obstacle, or falls off the edge of the map, the trial ends immediately. If both goals are collected, the trial ends early, several seconds after the second goal was collected. If both goals are not collected in time, the trial ends when the 30-second navigation timer is exhausted (which is again identified by a series of 3 chimes).

Following the end of each trial, an interstitial screen appears telling the participant that they can pull the trigger to proceed to the next trial. The study continues this way, until all trials are completed. At the end of the study, a screen appears providing feedback on the participant’s performance. This screen summarizes the number of goals collected, as well as the bonus compensation earned based on the participant’s performance. Base compensation was $20. Bonus was added as follows. 0-20% of all possible goals: $0; 21-40%: $1; 41-60%: $2; 61-80%: $3; 81-100%: $5. After having read this information, we helped the participant remove the head-mounted display.

Finally, participants respond to a brief survey using a laptop. When finished, we released the participant to receive their total compensation, and exit the study room.

#### Ethics statement

The experimental protocol was approved by the Committee for Protection of Human Subjects, under CPHS protocol number 2021-06-14394. Study protocol, as described above, included a written and verbal informed consent prior to the experiment. Study methods and provisional hypotheses were registered and can be viewed on the study’s OSF project page (https://osf.io/5tacn/). A video capture of the point of view of a sample participant is available as S1 Video.

### Analyses performed

#### Performance metric and other dependent variables

The simplest indicator of performance is the score obtained during the experiment (equal to the number of goals collected). Our analysis indicates that most participants devised planning strategies that were effective for the goal collection task. Mean score overall was 1.50 *±* 0.63. Mean score per map varied, and in some maps, no participants successfully collected both goals. Hence, to account for this diversity, we defined a more fine-grained metric of performance that also takes into account map difficulty, and spatial progress towards the second goal when not collected (see Table 1 for details).

**Table 1:**
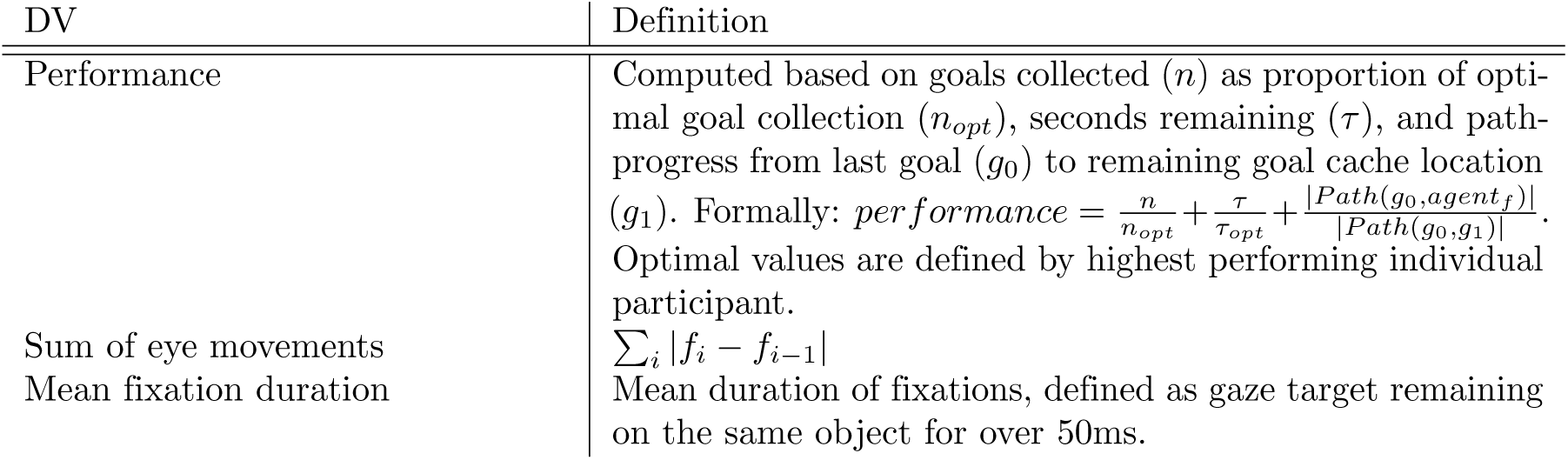
Description of key dependent variables.

Based on this latter performance metric, we split participants into three equally sized percentile-based performance group bins (bottom, mid, and top-performers), and compare bottom to top-performers. For these analyses, mid-level performers were ignored. The top-performing group show higher performance metrics on both 3 and 6-cache maps (bottom-performers: 3-cache map mean performance: 1.16, 6-cache map mean performance: 1.13. Top-performers: 3-cache map mean performance: 1.57, 6-cache map mean performance: 1.52).

Furthermore, to study the strategies followed by participants, we defined additional dependent variables, which consider the sum of eye movements, mean fixation duration, and the percent of forward and reverse ‘sweeps’ of eye movements across possible maze paths (see Table 1).

#### Salience of map features during planning

To identify which features of the map were considered to be more salient during the planning phase, we calculated an aggregate gaze heatmap for each map, across all participants, by summing tile-level gaze data during planning. Two methods of aggregation were explored: one based on fixation counts, and the other on total fixation duration. In both cases the resultant aggregate was standardized as an *N* -dimensional probability distribution (where *N* is the number of tiles in the base map).

Notably, these aggregate heatmaps are sequence agnostic, and therefore are interpreted as a map of static visual salience during planning.

Then, we mapped the flattened gaze distributions to two sets of features of the maps. The first set of features comprises the five tile types: water, muddy water, land, obstacle or goal. The second set of features corresponds to 9 feature maps that we identified, based on the spatial and informational geometry of each map. A description of each feature map tested is included in Table 2. Furthermore, Fig 2 provides example feature maps computed for map 2, as well as aggregated participant gaze on this map.

**Figure 2:**
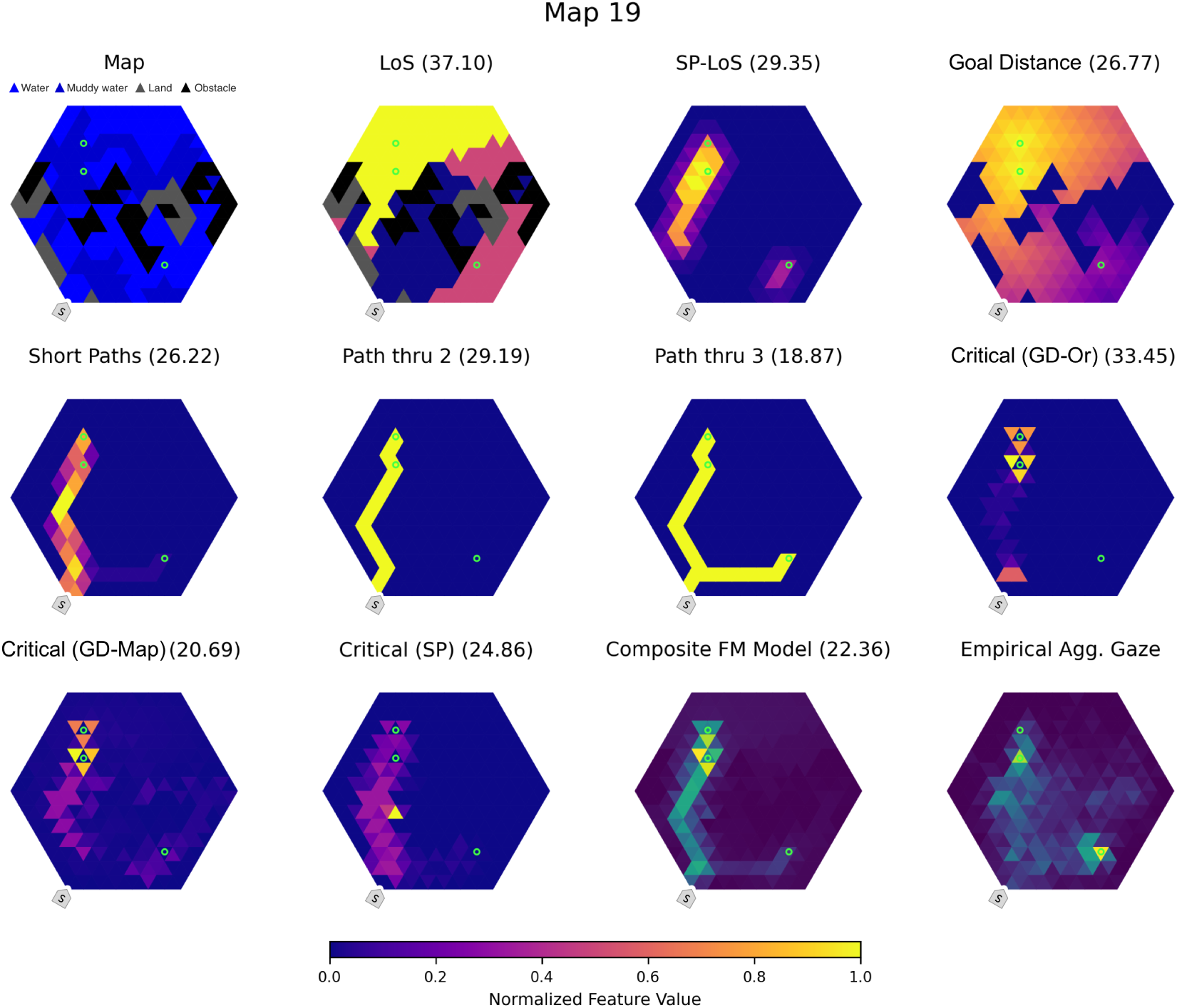
Sample feature and aggregated participant gaze for map 19. The first (top-left) figure shows map 19, in the format presented to participants. The subsequent 10 maps show different types of features, which comprise Line-of-Sight (“LoS”), Shortest-path Line-of-Sight (“SP-LoS”), distance to nearest goal (“Goal Distance”), Short Paths, Path through 2 goals, Path through 3 goals, Critical (via Goal-distance change at origin), Critical (via goal distance change map-wide), and Critical (via shortest path change). See Table 2 for details on each feature map. Note that the two critical feature maps identify bottleneck tiles such as small openings or possible shortcuts where inverting passability would enable or block a useful path. The mean of all feature maps is shown as the “Composite FM Model” in the second to last map. The last (bottom-right) figure shows the aggregated participants gaze for map 19. The numbers in parentheses above each map indicate the distributional distance between each feature-map and the aggregate gaze distribution using the 2D Euclidean earth mover distance (EMD).

**Table 2:**
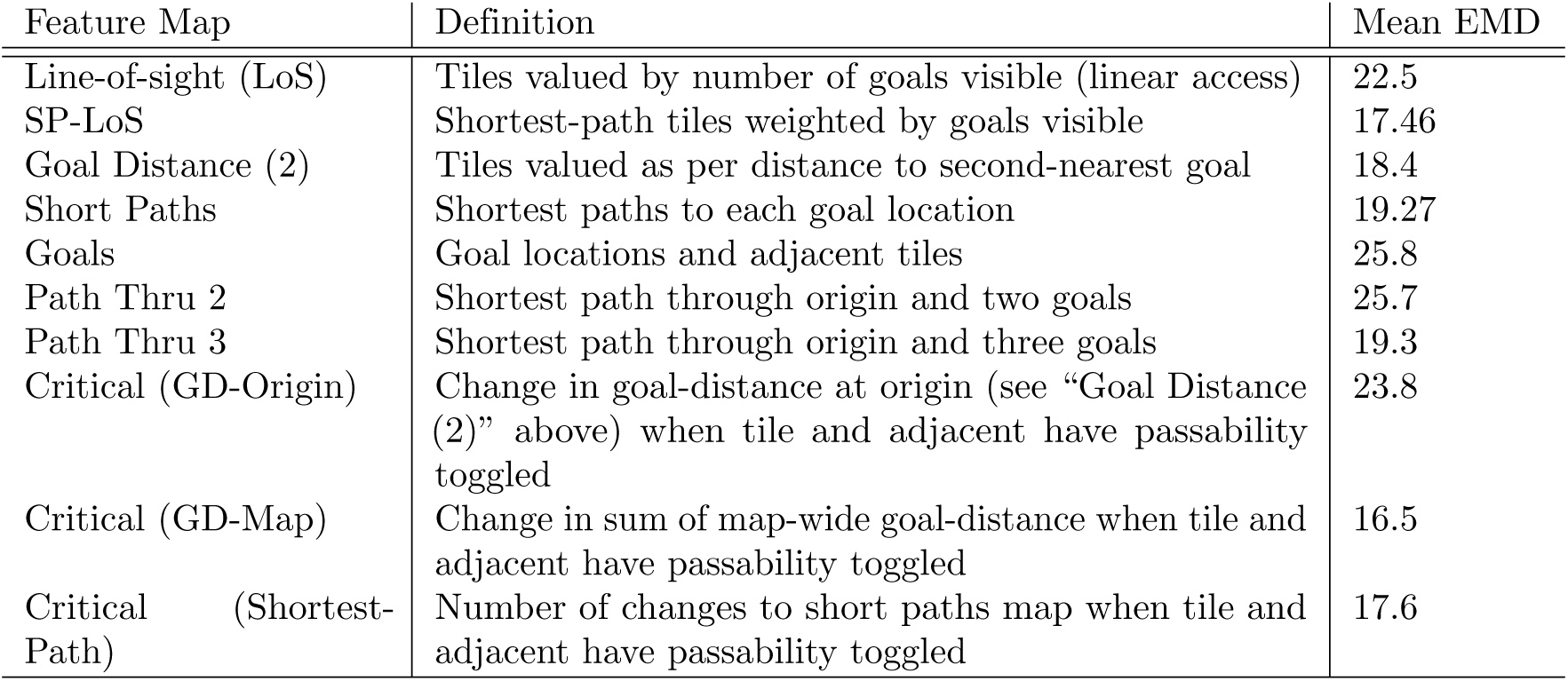
Description of feature maps used in prediction of planning gaze. The third column shows the mean cross-map Earth Mover Distance (EMD) between the feature map and the map-aggregate gaze heatmaps. The three “critical” feature maps are designed following analyses in Zhu et. al. which quantify the importance of a tile to the navigation task by comparing connectivity when the tile is passable versus when it is blocked [36]. In “Critical (GD-Origin)” we compare the impact on distance to goal at the origin. In “Critical (GD-Map)” we compare the total map-wide distance to the nearest goal. And finally, in “Critical (Shortest-Path)” we compare how much the shortest-path is impacted by each tile’s passability. See Fig 6 for visual comparison of feature-map EMDs.

At the map level, we compared the distributional distance between each feature-map and the aggregate gaze distribution using the 2D Euclidean earth mover distance (EMD), using the POT python optimal transport library [50]. The EMD provides a symmetrical measure of how far probability weights must be transported to match two distributions. This measure was chosen based on the intuition that gaze targets farther from predicted gaze density in 2D coordinates indicate a worse fit between the generated agent dynamics and empirical data. Feature maps with lower mean distance (across all maps) therefore are interpreted as supplying greater predictive power over preferred gaze locations.

#### Temporal gaze analysis and RQA

In addition to considering “static” gaze distribution, we extracted a number of temporal features from planning-time gaze data, including time-binned total saccadic distance (which approximates the total amount of eye movement during a trial), distance from origin, distance from nearest goal, multi-step sweeps, and a range of metrics derived from Recurrence Quantification Analysis (RQA).

RQA is an analytic method common in dynamical systems, and increasingly used to gain insights into temporal dynamics of eye tracking data [51]. We generated recurrence plots from the sequential tile fixation data during the planning phase, and extracted an array of common RQA metrics measuring attributes of the timeseries such as the likelihood of a fixation being recurrent (recurrence rate, *RR*), the longest recurrent subsequence (*L_max_*), and others. These were used as independent factors in the ANOVA analyses, to identify statistically significant differences correlated with both performance, as well as geometric map attributes.

#### Map analysis and classification

To address research questions about the influence of information and uncertainty-related attributes of particular maps on planning processes, we ran ANOVAs on a number of gaze-related attributes as dependent variables, against a list of map features which were hypothesized to be relevant to navigation planning. These included the length of the shortest path through 2, 3, and 4 goal cache locations (note that the last could only be computed for 6-cache maps), as well as a handful of metrics classified manually for each map which aimed to capture information access (e.g. where goal caches become visible in the landscape) and a simplified path-segment representation of each map. The coding process involved first producing a tree illustrating key segments and decision points, and then exhaustively considering goal cache observation outcomes (without knowledge of true goal locations).

We call the resultant representations “info-graphical,” given their tree like graphical structure, and sensitivity to visual information availability in the landscape. For an example, consider map 7 shown on the right side of Fig 3. At the origin (as with all maps by construction), no goal cache locations are visible. Participants must remember to navigate to the left, and by the time they reach about the midpoint of their journey to the narrow corridor at the top of the map, they will have visual access to two of the goal caches (square info nodes). At planning time, they contend with two outcomes: either one present goal is observed, or both are. Both outcomes result in full goal location knowledge, but in the first case (bottom fork in info-graph), after collecting the first goal they’ll need to continue around the low obstacle in the center of the map to collect the second. The target goal during this journey is fully observed, but some memory of the map structure is required to remember which way to navigate around the obstacle, and hence this latter segment is marked as memory-dependent (three hatch marks). Alternatively, if both closer goals are present at the info node, they can continue around and collect both with no significant memory dependence (un-hatched edge).

**Figure 3:**
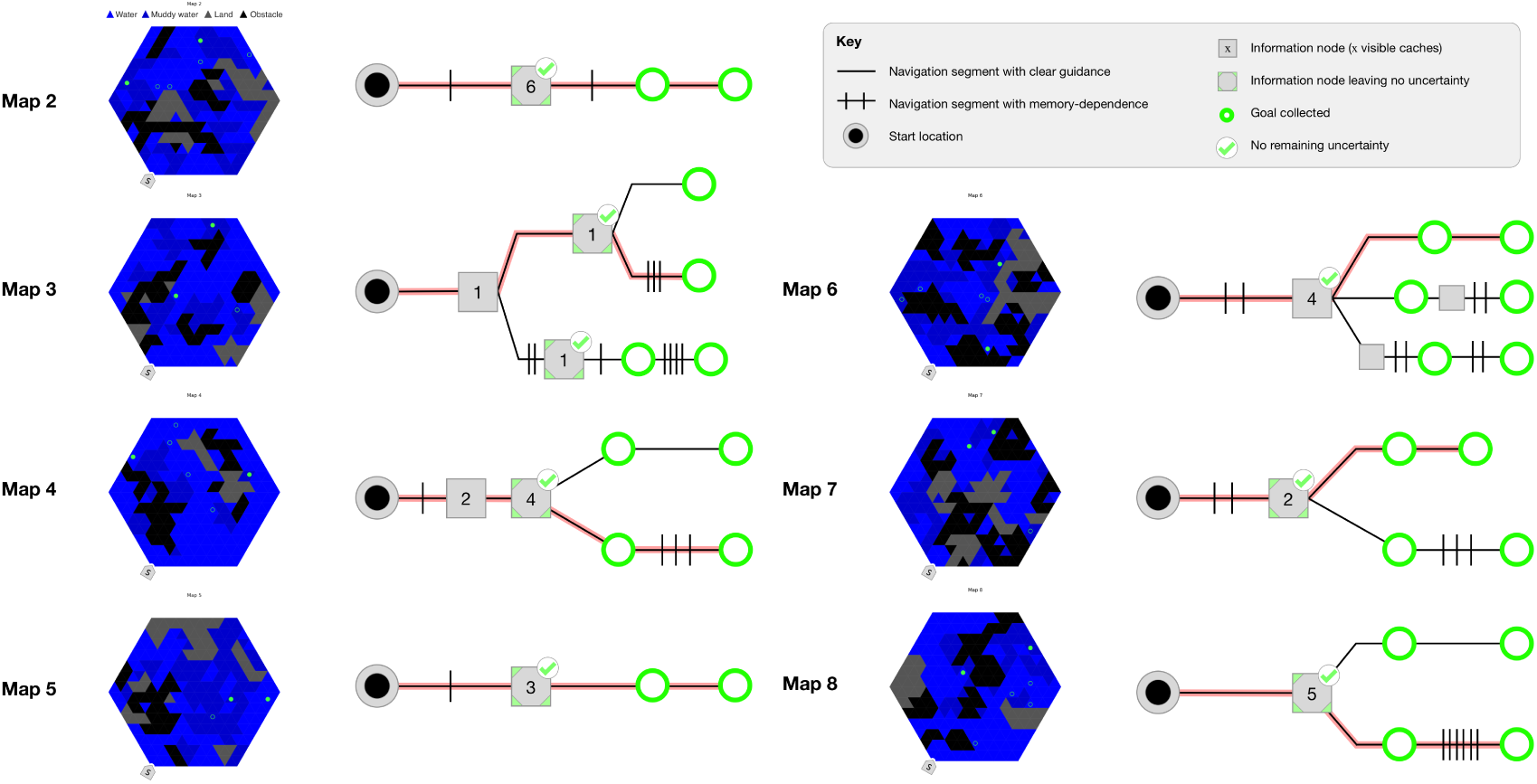
Map info-graphical classification for selected maps. See key for symbol semantics. Red line indicates expected path through graphical structure according to true goal locations.

After producing info-graphical representations for all maps, these were used to calculate a number of key metrics, including: branching factor, number of forks, steps to certainty, and expected number of memory-dependent segments. See Fig 3 for sample info-graphical representations of selected maps, and Table 3 for a full list of attributes computed.

**Table 3:** Key map attributes analyzed.

| Attribute | Definition |
| --- | --- |
| Branching factor | Average number of children of each node in the info-graphical path tree. |
| Number of forks | Count of forks in which a node splits into two or more children in the info-graphical path tree. |
| Number of information nodes | Number of informational nodes (represented as squares) in which goal-cache locations are hypothetically visible at navigation-time. |
| Steps to certainty | Expected number of nodes at which point all goal-cache presence statuses will have been observed. |
| Expected memory-dependent segments (planning) | Expected number of tree segments that require some memory of map structure to successfully navigate (indicated by hash marks through path segment). |
| Expected memory-dependent segments navigated | Expected number of tree segments to be navigated conditioned on true goal locations. |

A sample in-map view of an example information node that results in no remaining uncertainty can be seen in the sample “Participant POV” video (S1 Video). At 29 seconds into this screen recording, showing Map 6, the participant observes 1 goal among the 3 central goal caches, and is in position to confirm whether the final goal is to the right or left. This is represented by the 4-cache information node in Map 6 of Fig 3. We note that in this case the participant continues to the observed goal without confirming the second goal location.

## Results

### Trends in gaze dynamics through planning

The first question that we address in this study is whether gaze dynamics show signatures of hierarchical planning and subgoaling and whether there are differences between top and bottom performers. For this, we focus on temporal gaze dynamics prior to movement.

Fig 4 shows that consistent trends in gaze dynamics can be found across the 15-second planning phase. Total saccade distance decreases while fixation duration increases. Fixation targets converge to tiles closer to the path ultimately taken, with minimum distance seen at the end of planning (see also S1 Fig) for gaze dynamics in some example maps.

**Figure 4:**
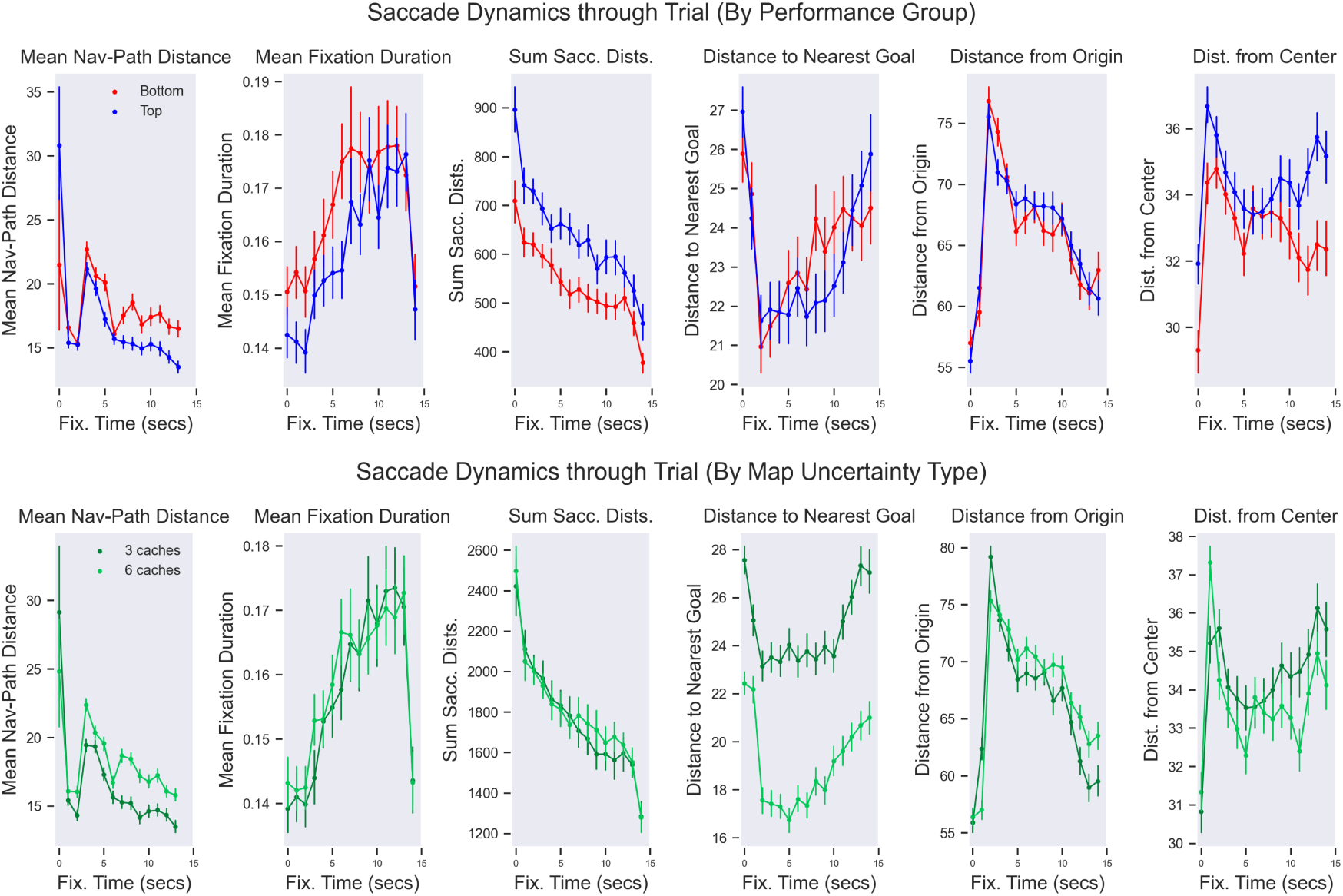
Trends in eye movement dynamics through planning phase. The six plots show the distance between gaze and navigation path, the saccade distance, saccade duration, distance from nearest goal, distance from origin, and distance from center. Top panel: comparison of top and bottom performers. Bottom panel: comparison of 3 and 6-goal maps.

Additionally, a U-shaped trend is observed in distance from fixated tiles to the nearest goal location; tiles close to goals are more frequently fixated in the middle of the planning phase, while more distal tiles are viewed at the start and end of planning. These dynamics are largely consistent across top and bottom performers (top panels) and across 3 and 6-goal maps (bottom panels), but among top performers, gaze exhibits lower fixation durations and greater movement distances.

Finally, when gaze metrics were analyzed against map attributes, results show a consistent relationship between key map info-graphical attributes and eye movements. Specifically, maps with more forks, a higher branch-factor, and more memory-dependent segments are associated with longer eye movements and reduced fixation duration during planning. Furthermore, we found associations between map attributes and planning-time gaze recurrence metrics. The map branch factor, number of memory-dependent segments, and number of forks were associated with lower recurrence rate (RR). These results may indicate wider novel exploration (which therefore exhibits fewer repetitions during search) in more complex maps. See Table 4 for a summary of ANOVA results.

**Table 4:** Summary of ANOVA results comparing gaze dynamics with map-level info-graphical metrics. “Dir” indicates whether the relationship is positive (+) or negative (-). Planning RR is gaze recurrence rate during the planning phase. Significant results (with *p ≤* 0.05) are marked in bold.

| DV | Factor | DF | $F$ | $p$ | $\eta_p^2$ | Dir. |
| --- | --- | --- | --- | --- | --- | --- |
| Sum of eye mov. | Branch factor | $F(2, 66)$ | 9.86 | **< .001** | .011 | + |
| Sum of eye mov. | No. forks | $F(2, 66)$ | 13.3 | **< .001** | .013 | + |
| Sum of eye mov. | No. memory-dep. segs. | $F(1, 33)$ | 4.9 | **.033** | .004 | + |
| Mean fix. dur. | Branch factor | $F(2, 66)$ | 4.99 | **.010** | .006 | - |
| Mean fix. dur. | No. forks | $F(2, 66)$ | 6.47 | **.003** | .006 | - |
| Mean fix. dur. | No. memory-dep. segs. | $F(1, 33)$ | 10.2 | **.003** | .004 | - |
| Planning RR | No. forks | $F(2, 66)$ | 8.2 | **< .001** | .027 | - |
| Planning RR | No. memory-dep. segs. | $F(1, 33)$ | 22.6 | **< .001** | .032 | - |
| Planning RR | Branch factor | $F(2, 66)$ | 4.1 | **.020** | .015 | - |

### Gaze distribution and the salience of map attributes

The second question that we address in this study is what features of the map are considered more salient during planning — either because they are candidate goal or subgoal locations or because they impair planning (e.g., obstacles that require detours) — and whether there are differences between top and bottom performers.

We considered the visual salience of map attributes through a static analysis of aggregate (cross-participant) gaze on each map. Fig 5 shows the fixation counts and duration in relation to the five different tile types (water, muddy water, land, obstacle or goal tiles). The value of zero in the plot indicates expected fixation duration and time if visual attention was perfectly uniform for a given map. Our analysis indicates that both goal locations and muddy water are disproportionately fixated at planning time. Goal locations are obviously salient for the task, whereas muddy water (and especially unavoidable muddy water along a chosen path) might require more careful consideration.

**Figure 5:**
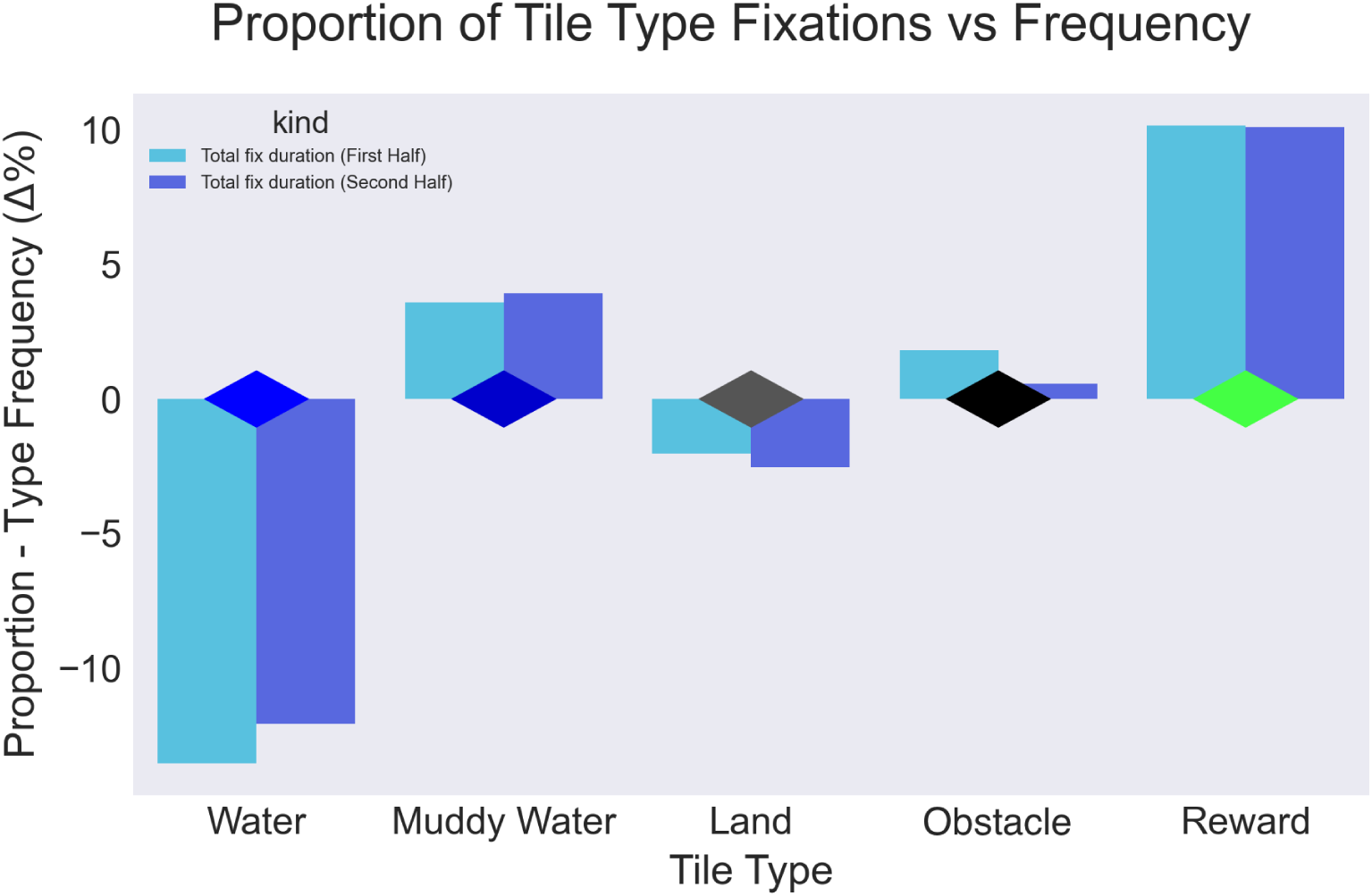
Tile type fixation proportionality. The value of zero in the plot indicates expected fixation duration and time if visual attention was perfectly uniform for a given map.

Furthermore, we found that regular water tiles were under-fixated by both metrics, likely due to regions of each map easily identified as not relevant to the navigation task, and therefore pruned early on. We also found that fixations on muddy water and goal tiles were even more disproportionately favored on 6-cache as compared to 3-cache maps. Finally, we found decreasing attention on obstacles in the second half of the planning phase.

When comparing map attribute salience by performance group, we find a number of differences that may suggest attention-correlated sub-goaling strategies that were useful to performance. Bottom performers fixated longer on obstacle tiles as compared with top performers, especially during the second half of planning. We also find that bottom performers spent more time at the beginning of planning attending to “first fork” regions. First forks were defined by selecting a region around the first tile where any optimal path to goal diverged. These locations are often readily apparent as early points where a navigation decision will need to be made. Likewise, during the first 2 seconds of planning, top performers more quickly identified another key point of salience: the actual first goal they would collect.

We also analyzed another plausible sub-goal type critical to ambiguity reduction during navigation. The region of “first certainty” is defined as the closest location from which the exact position of all goals will be determinable. More specifically, first certainty is a line-of-sight based analysis that follows all paths to goals, and determines the earliest location where *n −* 1 goal cache locations will be visibly accessible. From this point on the map, it is possible to infer the true location of each goal, by inspecting all (or all but one) goal caches. Top performers were found to attend to this region sooner than bottom performers on both 3 and 6-cache maps.

Fig 6 shows the fixation counts and duration in relation to the static feature-maps described in the Methods. We found that the feature maps were able to predict planning-time gaze in aggregate (across participants) significantly better than a uniform baseline. Feature-maps based on tile criticality—that is, the sensitivity of navigation paths and goal distances to the passability of particular locations—had the lowest average EMD, implying they offer the highest predictive value (see Table 2 and Fig 6). See also S2 Fig (and continuations in S3 Fig, S4 Fig, and S5 Fig) for illustrative examples of how feature maps reflect gaze allocation.

**Figure 6:**
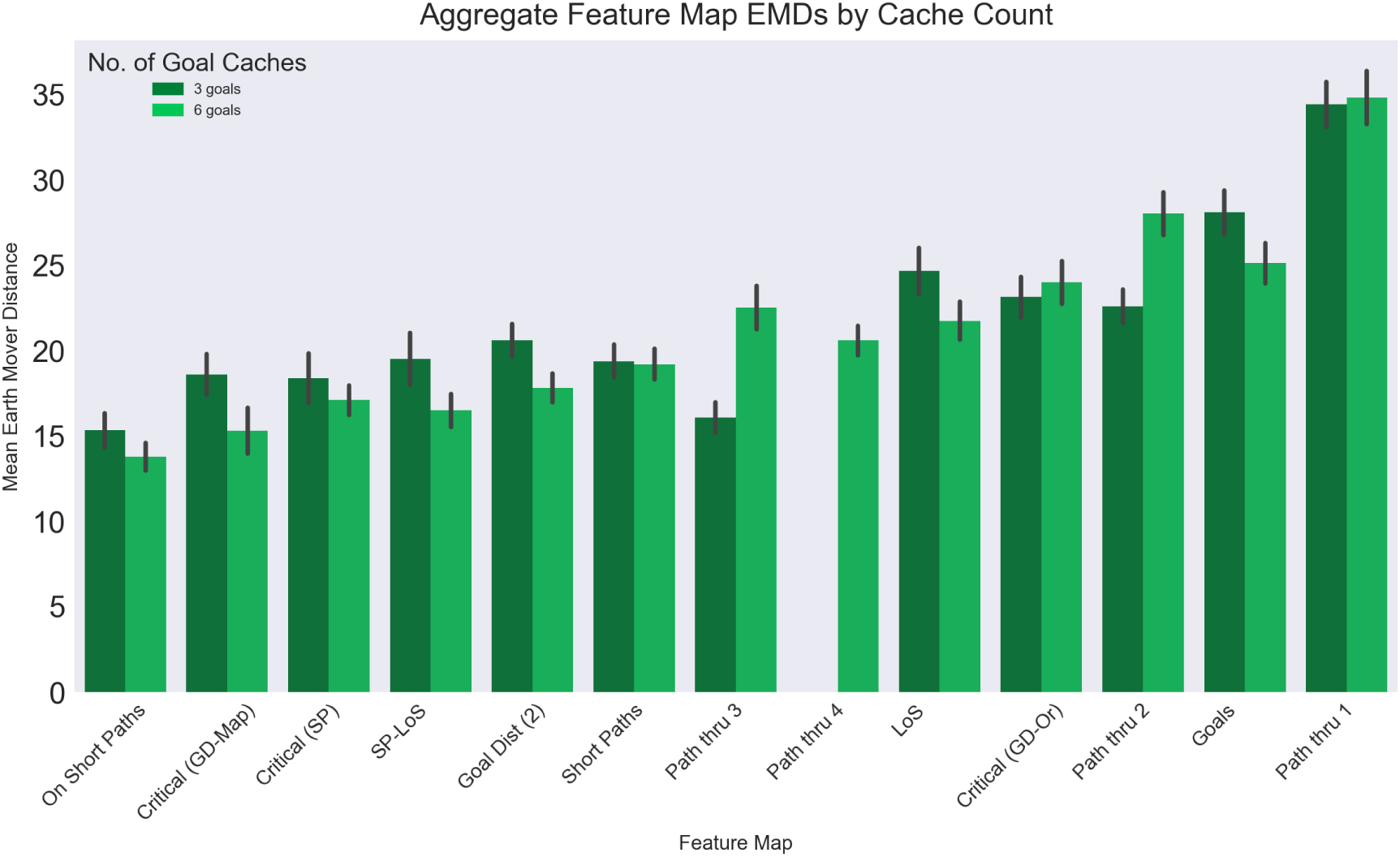
Feature-map predictors of aggregate gaze maps. Lower EMDs correspond to greater distributional similarity between feature map and aggregated participant gaze. Bars are plotted in ascending order from lower to higher distances.

### Relation between path planning dynamics and performance

The third question that we ask is whether and how the planning strategy of participants prior to movement influences their subsequent performance. We address this question by considering potential differences between path planning depth and initial movements during planning of top-and bottom-performers.

#### Relation between path planning depth and performance

We first ask whether top- and bottom-performers differ in how much they plan ahead during the planning phase (i.e., “planning depth” [52]).

Included in the feature-map analysis were maps generated by finding the shortest path through both the origin and either 1, 2, 3 or 4 goal locations (SP-1, SP-2, SP-3, and SP-4 respectively). Using these maps, we inferred each participant’s “planning depth” – or the putative depth of the plans that participants developed during the planning phase – on each trial, by choosing the SP-depth map with the minimum distance (EMD) to trial-level planning gaze.

We therefore analyzed the relationships between these four levels of planning depth (SP-1 to SP-4) and both performance and various map attributes. As shown in Fig 7, in three of the info-graphical attributes most directly correlated with map difficulty (number of forks, branch factor, and number of informational nodes), we see a relative agnosticism to depth class for simpler maps (e.g. those with less than 2 forks, or less than 3 info nodes). However, in the highest complexity group, a stronger performance correlation appears. Participants with planning depth most similar to SP-3 performed best on maps with the largest number of forks, while SP-4 participants performed best on maps with the largest number of info nodes, and largest branching factor.

**Figure 7:**
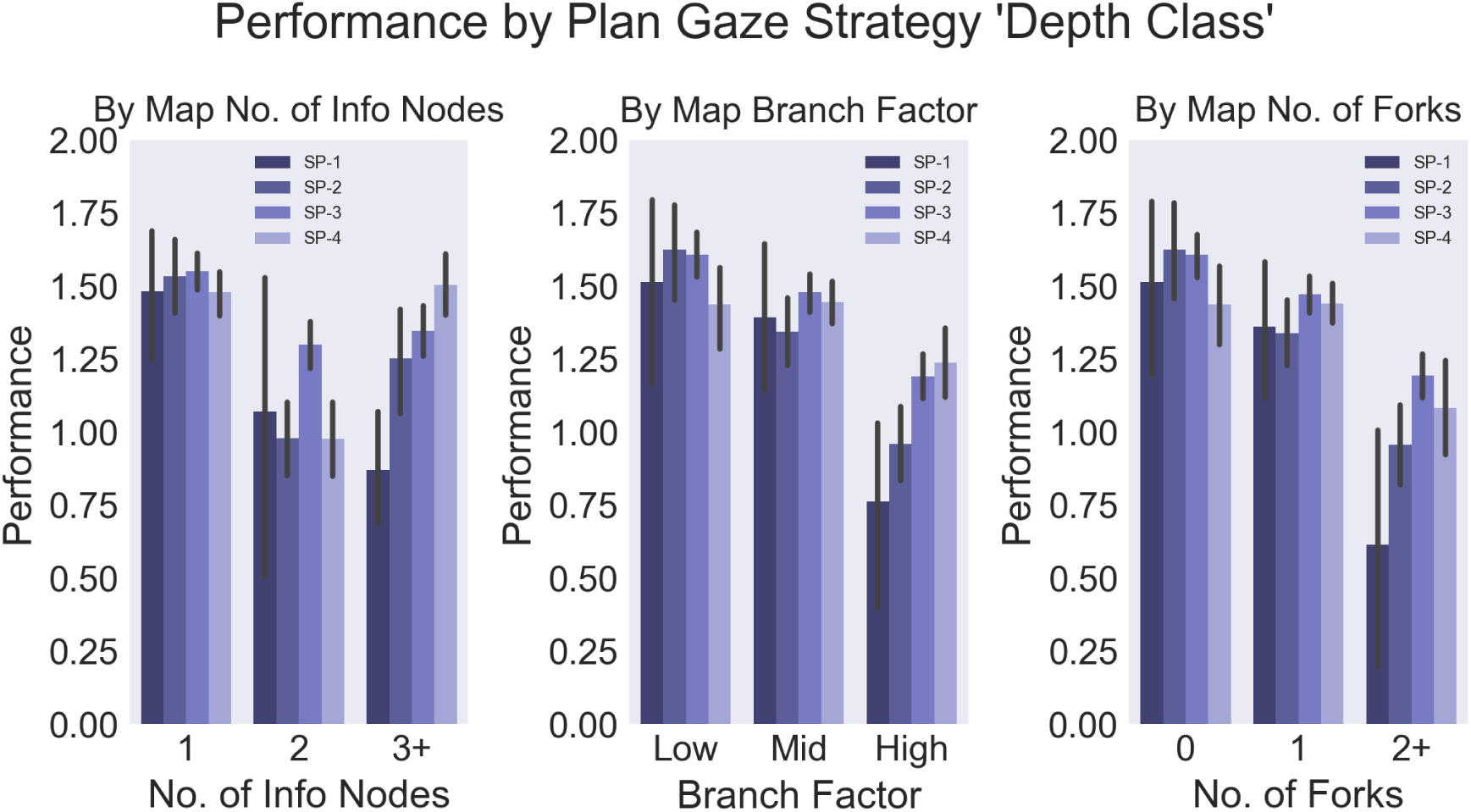
Performance and planning depth. Trial-level performance comparison showing the relationship between planning depth (as inferred from planning time gaze distance from shortest-paths through 1, 2, 3, and 4 goal caches), and info-graphical map attributes. A positive, nearly monotonic, relationship is seen in the most complex maps (as determined by number of forks, branch factor, and number of info nodes) between planning depth and performance.

When bucketed by performance group, we find SP-3 and SP-4 depth classes were more common among top-performers. The SP-3 class was seen in 47.1% of bottom, as compared with 51.0% of top performers. The SP-4 class was seen in 26.7% of bottom, as compared with 30.1% of top performers.

Selected animations showing both gaze and navigation trajectory through each trial, are available as Supporting Information (see S2 Video, S3 Video, and S4 Video). All map animations can be viewed on the OSF project page in the “Aggregate Gaze Animations” directory (https://osf.io/5tacn/). In these animations, full trial dynamics can be seen, aggregated by top and bottom performers (left and right maps). Two different hysteresis rates are shown (top: *λ* = 0.93, fast decay, and bottom: *λ* = 1.0, no decay) to better visualize the evolution of each gaze distribution over time. Several of the above noted planning-performance relationships can be observed when viewing individual trial timeseries, e.g. in Map 20 (S4 Video), top performer gaze reaches the 3 concentrated goal caches towards the right of the map earlier and more densely than bottom performers.

#### Relation between arm movement during planning, initial heading, and performance

Finally, we ask whether arm movements during planning are predictive of some of the very first decisions made—which direction to begin navigating—in bottom- and top-performers. For this analysis, we defined initial navigation direction based on the position of the agent in relation to the map center-line (“left” or “right”), 1 second after the beginning of navigation^1^.

We find that, indeed, arm movements are directly correlated with initial heading, implying that the initial movement direction can be read out from the position of the yaw controller prior to the beginning of movement. This trend might reflect planning and preparatory processes leaking into the execution phase or a way planners offload part of their cognitive processing to physical activity [53–56]. Notably, while this trend is very clear in top-performers, it is much less pronounced in bottom-performers, in which the average position of the yaw controller remains very close to the middle line. See controller yaw traces, segmented by initial navigation direction (left and right panels segment trials by initial navigation to the left or right, respectively), and performance group (blue and red lines show data from top and bottom-performers, respectively), in Fig 8. The higher-amplitude movements seen at the beginning of the navigation phase (top of plot) illustrate the larger movements required to control agent rotation as navigation began. As expected, this portion exhibits a heavy directional bias due to the definition of the “initial navigation direction” variable used for segmentation.

**Figure 8:**
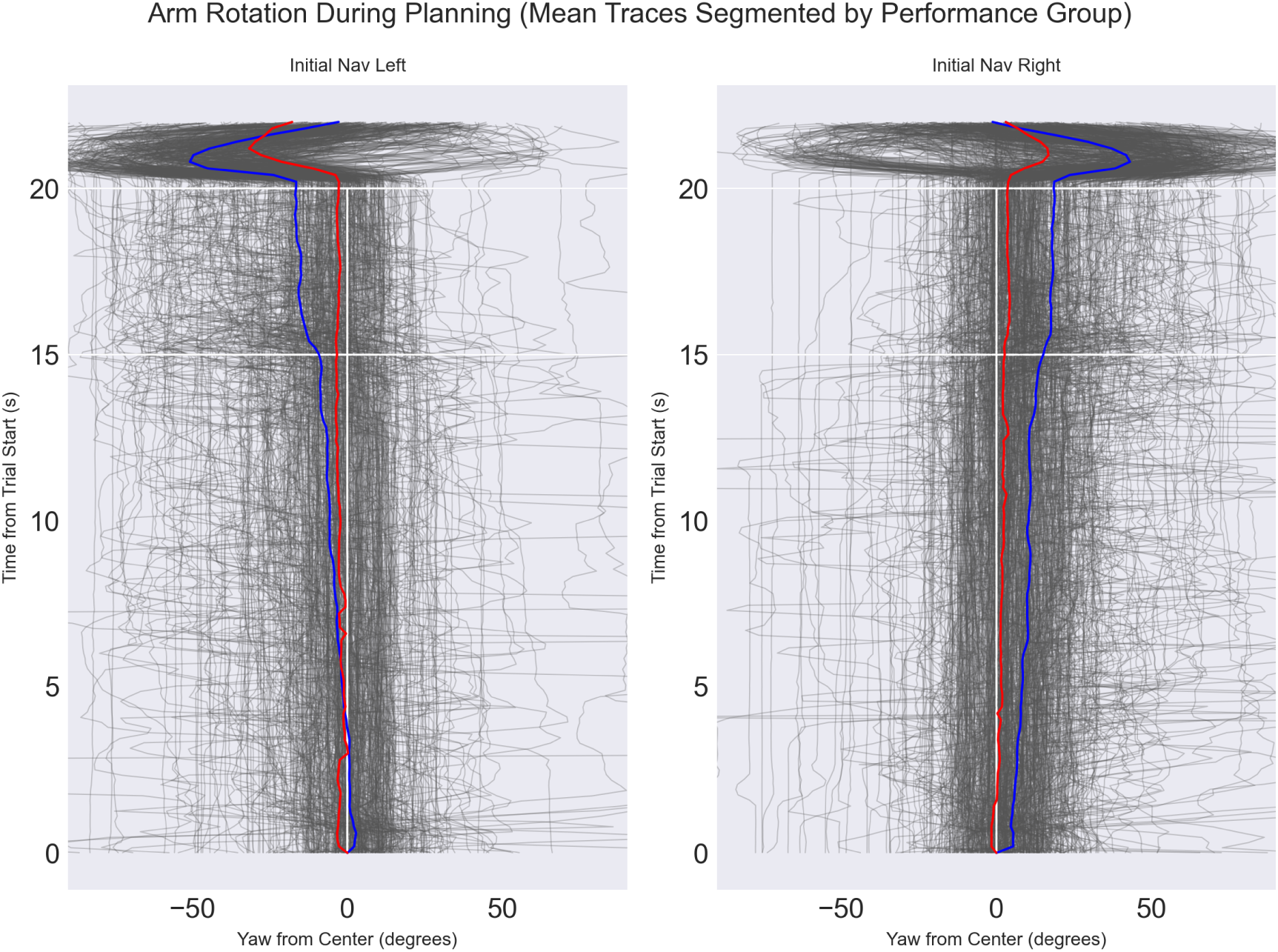
Arm movement traces as measured by controller yaw. Traces are segmented by initial navigation direction (left and right panels correspond to initial navigation to the left and right, respectively). Time from trial start progresses vertically from bottom. Navigation direction groups are defined as the sign of the offset from center-line 1000ms after navigation start. The blue and red lines show mean across trial-level traces among top and bottom-performers, respectively. Planning phase (bottom 15 seconds), transition (following 5 seconds), and first 5 seconds of navigation phase are shown, separated by white lines.

## Discussion

In this human behavioral study, participants were asked to view an overview of a landscape for 15 seconds, prior to immersively navigating through it to find and collect up to two goals. Because the number of possible goal locations (“caches”) exceeds the number of true goals, successful participants had to develop a navigation plan robust to this uncertainty. A primary motivation in this design is to better understand and characterize the dynamics of the internal representations developed during planning to guide future movement, and the handling of uncertainty during this process. However, these representations were not, themselves, directly measurable. Instead, we analyzed eye and arm movements collected during the planning phase of the task, which putatively reflect some dimensions of the cognitive processes of interest, and admit quantitative comparison with geometric, visual, and informational features of the landscapes studied.

This work pursued three aims: determining whether gaze dynamics prior to movement show signatures of hierarchical planning and subgoaling, identifying the most relevant features for subgoal selection, and assessing which strategic aspects of planning distinguish top- from bottom-performers.

Our spatial and temporal analyses, as well as correlations between gaze patterns prior to navigation and quantitative map-level attributes and task performance metrics, show signatures of a planning process that is both hierarchical and bi-directional. These include a decreasing trend seen in gaze distance from origin, and a broad to narrow shift (with reducing saccade distances and longer fixation durations) as plans are established. These signatures of hierarchical planning likely aid participants in efficiently transforming rich spatial information into a simpler, contingency-aware, plan of action. Furthermore, in line with prior work, we identified “critical tiles” to which landscape connectivity is most sensitive as the features that best predict visual attention. Finally, performance-group-level analyses show that the most effective participants (top-performers) navigated paths closer to regions of the map visually attended during planning, exhibited more frequent eye movements with shorter fixations, as well as earlier attention to ambiguity-reducing sub-goals. Top performers always showed greater focus on more peripheral tiles and deeper planning depth, and leveraged offloading of initial navigation direction through orientation of the arm. Below we discuss these results in more detail.

### Gaze dynamics prior to navigation show signatures of hierarchical planning

In a temporal analysis of gaze dynamics, we find consistent trends indicative of hierarchical planning across the planning phase (see Fig 4). Total saccade distance decreases while (in line with this result) fixation duration increases. Fixation targets converge to tiles closer to the path ultimately taken, with minimum distance seen at the end of planning. A U-shaped trend is observed in distance from fixated tiles to the nearest goal location; tiles close to goals are more frequently fixated in the middle of the planning phase, while more distal tiles are viewed at the start and end of planning. Finally, gaze distance from origin shows a quick radiation outwards in the first several seconds, followed by a consistent negative trend for the remainder of the planning phase.

Together, these results illustrate that temporal gaze dynamics start broad and then spatially narrow as plans are established. Visual exploration first favors proximal locations, and is initially wide and fast, exhibiting saccades that move between easily-identified salient locations such as goal caches. Gaze then narrows (in terms of spatial spread), and deepens (seen as longer fixation durations) through the planning phase as a navigation plan is converged upon. These gaze dynamics could reflect a covert planning process that exhibits a hierarchical nature. Initial exploration of key landmarks may help to establish navigational affordances based on abstract map connectivity. Then, participants might establish a plan hierarchically, by enhancing an initially abstract path through high level navigation goals with lower level details of plausible routes between them. These results are consistent with previous findings on hierarchical organization of action execution [25–29, 41] and shed light on the temporal and visual dynamics enabling hierarchical plans to be established before movement.

### Salient tiles and critical regions predict attention during planning

Findings from map-based spatial prediction of gaze preferences suggest that visual search strategies are highly sensitive to not only goal locations and other gross aspects of spatial geometry, but also to more subtle features and aspects of graphical connectivity and information geometry (e.g. goal presence information available at a particular tile).

Overall, we find that goal locations and muddy water were most disproportionately fixated. We also observe increasing attention on water, and decreasing attention on obstacles in the second half of the planning phase (see Fig 5). This finding may indicate that early saccades to obstacles support the development of a simplified graphical representation of map connectivity (and information access), while later gaze sequences support evaluation of candidate paths through the structure identified. These results are in agreement with other studies showing that people form simplified problem representations to plan [15] and that cognitive task representations can be distorted, removing task-irrelevant dimensions [57].

Gaze prediction analysis using static geometry-based feature-maps enabled a more nuanced analysis of the types of features that attracted visual attention. Feature-maps were effective predictors of planning-time gaze in aggregate (across participants). Comparison of predictive power for each feature-map analyzed is suggestive of the landscape features that most consistently attracted planning-time visual attention (see Table 2 and Fig 6). Feature-maps based on tile criticality, which identify regions for which a change to connectivity (i.e. switching a passable region for an obstacle) has particularly outsized effects on key navigation considerations such as the lengths of efficient paths (Critical Shortest-Path), or mean goal distance map-wide (Critical GD-Map), were especially useful predictors. This finding is consistent with results related to edge “toggling” in [36] and shows that it generalizes to tasks requiring participants to develop robust navigation plans.

### Top-performers show distinctive planning strategies along multiple dimensions

Our results show that several strategic aspects of planning distinguish performance groups. First, we find that among top performers, gaze exhibits lower fixation durations and greater movement distances. These attributes are correlated with a higher throughput of information processing at planning time and might reflect a more efficient allocation of limited resources [45].

Second, we find that attention in some key map regions, especially early attention in the first few seconds of planning, differ by performance groups. A bias in attention towards the first fork is seen in bottom-performers, which may indicate that these more superficial or readily apparent map features attracted more attention for this group, while top-performers may have more quickly moved to subtler decision points such as deeper cache-location dependent forks, or more distant clusters of plausible goals. Additionally, we find that “first certainty” sub-goal locations were attended earlier by top performers, possibly reflecting a greater attention to epistemically critical sub-goals conveying full-map information (the location of all goals), but which require a richer (perhaps simulative) forecast of visual and informational dynamics at navigation time. Together, we interpret these findings as indicating that different performance groups’ attention was attracted to different kinds of sub-goals, and at different times. First forks are readily apparent from local (near origin) map structure, whereas identifying which goal is most plausible to capture, or which locations will provide maximal ambiguity reduction, requires deeper consideration, and perhaps more robust simulation. This may explain why attention in these regions better differentiates skill and performance.

Third, we find that deeper planning gaze predicts success on high-uncertainty maps. Planning depth results showed that the SP-3 map was the most predictive of aggregate gaze across participants (on both 3- and 6-cache maps), indicating a possible balance of (cognitively costly) planning depth and the risk of planned caches being empty. Performance correlations with gaze depth-class (as determined by similarity with “SP-X” feature maps) indicate that some map attributes rendered deeper plans particularly helpful. In particular, planning gaze similar to SP-3 was associated with highest performance on maps with at least one fork. Similarly, maps with fewer information-nodes (locations at which goal existence information would be obtained) saw greatest performance when gaze resembled SP-3. In contrast, SP-4 performed best in the most epistemically complex maps with the largest number (3 or more) of informational nodes. See Fig 7 for comparisons, and the Methods for details on the determination of map classification attributes. Another perspective on this result can also be seen in the performance-binned gaze preferences in Fig , which show a bias for more peripheral, and later path segments among the top performing group. We interpret these findings as evidence that deeper plans predicted performance, but only on more challenging maps where multiple branches, or multiple stages of information access, required the development of a more robust plan (one with visits to more goal caches). On the other hand, a greedier (and riskier) strategy was satisfactory on maps with simpler information geometry and less path uncertainty.

Finally, we find that the behavior of top-performers before navigation better predicts their initial navigation direction compared to bottom-performers. The fact that it is possible to decode initial navigation direction before navigation starts reflects the fact that there is not a clear cut separation between the planning and execution phases, but rather some aspects of the plan leak into the overt execution [56, 58–60]. However, this trend is much more prominent in top-performers, which may suggest that they *exploited* the continuous nature of planning and execution in the task at head. These findings may be signatures of offloading part of the decision memory to physical state, potentially alleviating cognitive burden [53–55, 61].

### Limitations & future work

In this study, uncertainty was primarily manipulated by requiring participants plan with incomplete information about true goal locations. While this paradigm corresponds with certain naturalistic settings where rewarding states often fall within a set of discrete locations, other types of uncertainty are frequent in real-world situations requiring planning and navigation. As one example, path connectivity may differ from maps or other representations available when a route is being selected, e.g. due to unforeseen obstacles, the presence and influence of other individuals (e.g. crowds that slow one’s progress), and other modifications to the environment (e.g. a transit line discovered to be down for maintenance). These environmental uncertainties likely require even more robust contingency planning, and perhaps prompt the development of different internal representations better suited to more continuous and dynamic changes. Such a design would likely limit the efficacy of heuristic pathfinding strategies such as “find a short path through 4 goal caches,” likely used by some participants in this study, since path length would itself be unpredictable during planning. Eye movements and other biometric data would likely shed light on both correspondence and divergence in the planning processes used by individuals encountering these other types of uncertainty.

Similarly, while the plan-then-execute paradigm employed here reflects common daily scenarios where planning may progress prior to direct engagement in the environment, as discussed in [58], more embodied planning and action scenarios require the constant interplay of perception and action as the state of an individual with respect to their environment continuously evolves. The meta-cognitive and resource-rational aspects of how to coordinate movement with sensory sampling, and when and how to leverage deeper simulations for more distal planning processes, even during ongoing activity, is not well understood; these questions offer many rich avenues for further study. While analyses comparing planning gaze to gaze at time of navigation were out of scope for this analysis, an exploration of this relationship, especially as it relates to expertise and group-level performance differences, is an exciting avenue for future research. Aggregate gaze trajectories were collected and visualized for each map, and include gaze through the full trial (spanning both planning and navigation time). These can be viewed in the sample Supplementary Information (e.g. S2 Video) as well as in the comprehensive video list for all maps on the OSF project site.

## Conclusion

This work presented a virtual-reality based behavioral experiment which collected eye movements during a pre-navigation phase, with the aim of capturing aspects of covert planning processes related to pathfinding through a landscape with uncertain goal locations. Taken together, our results demonstrate the way gaze, and other biometric signals, can uncover hierarchical attributes of planning dynamics through time. Furthermore, we show that the nature of the hierarchy built (e.g. the set of sub-goals chosen as its nodes) is skill-dependent — highlighting the capability of top-performers to allocate their cognitive resources to task-relevant features and to form deeper plans. These results showcase the importance of devising effective planning and active exploration strategies to solve challenging navigation tasks that defy exhaustive search.

## Data availability

The data and materials for all experiments are available at https://osf.io/5tacn/.

## Supporting information

**S1 Fig.**
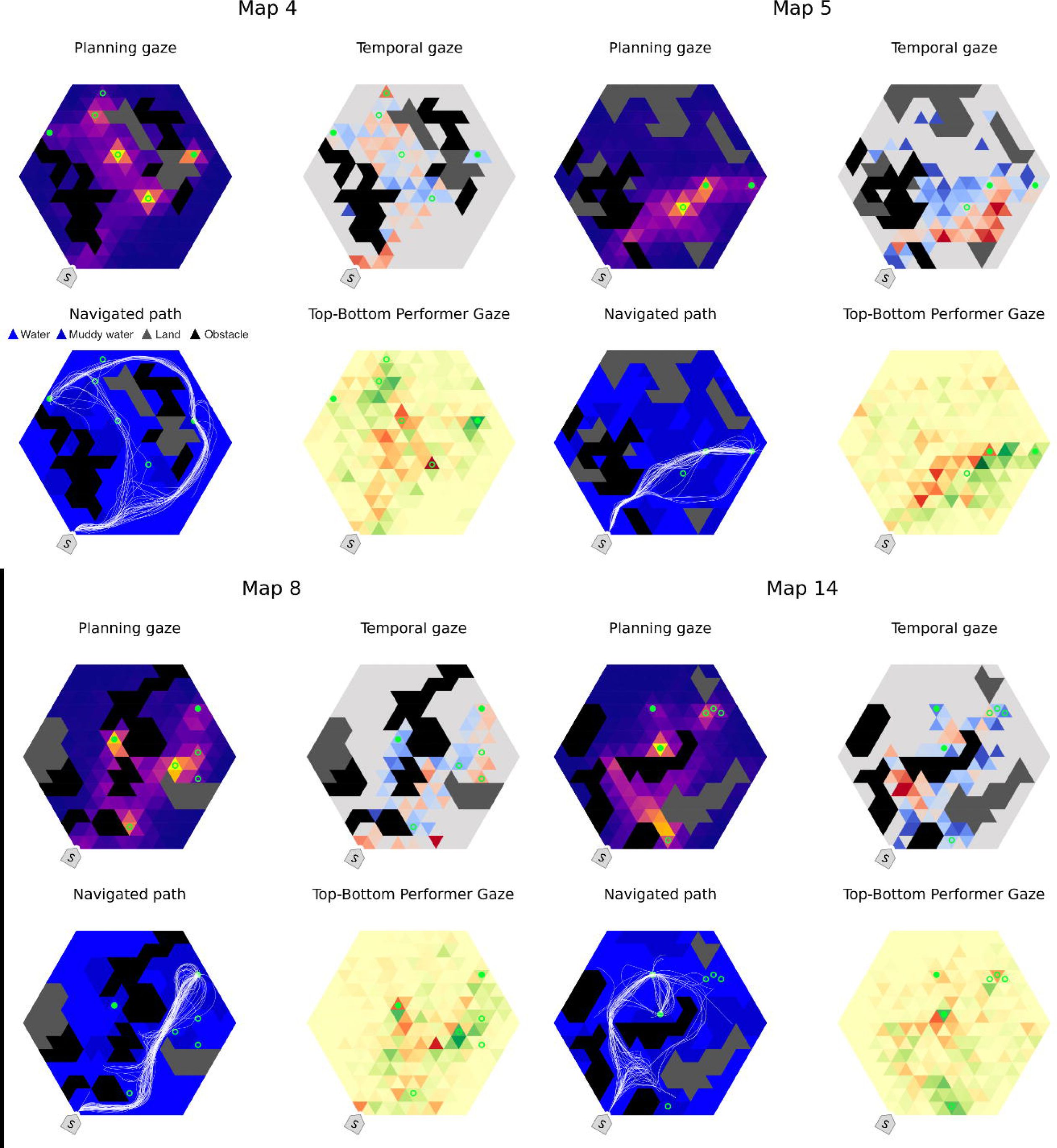
Sample gaze and navigation analyses for four maps. For each map, the following analyses are visualized: aggregated planning gaze (upper left), temporal gaze shift (upper right; blue tiles viewed earlier during planning, red later), path navigation trajectories (lower left), and performance-group differences in gaze density (lower right; green tiles show greater density among the top-performing group, and red tiles are those with greater density among the bottom-performing group.)

**S2 Fig.**
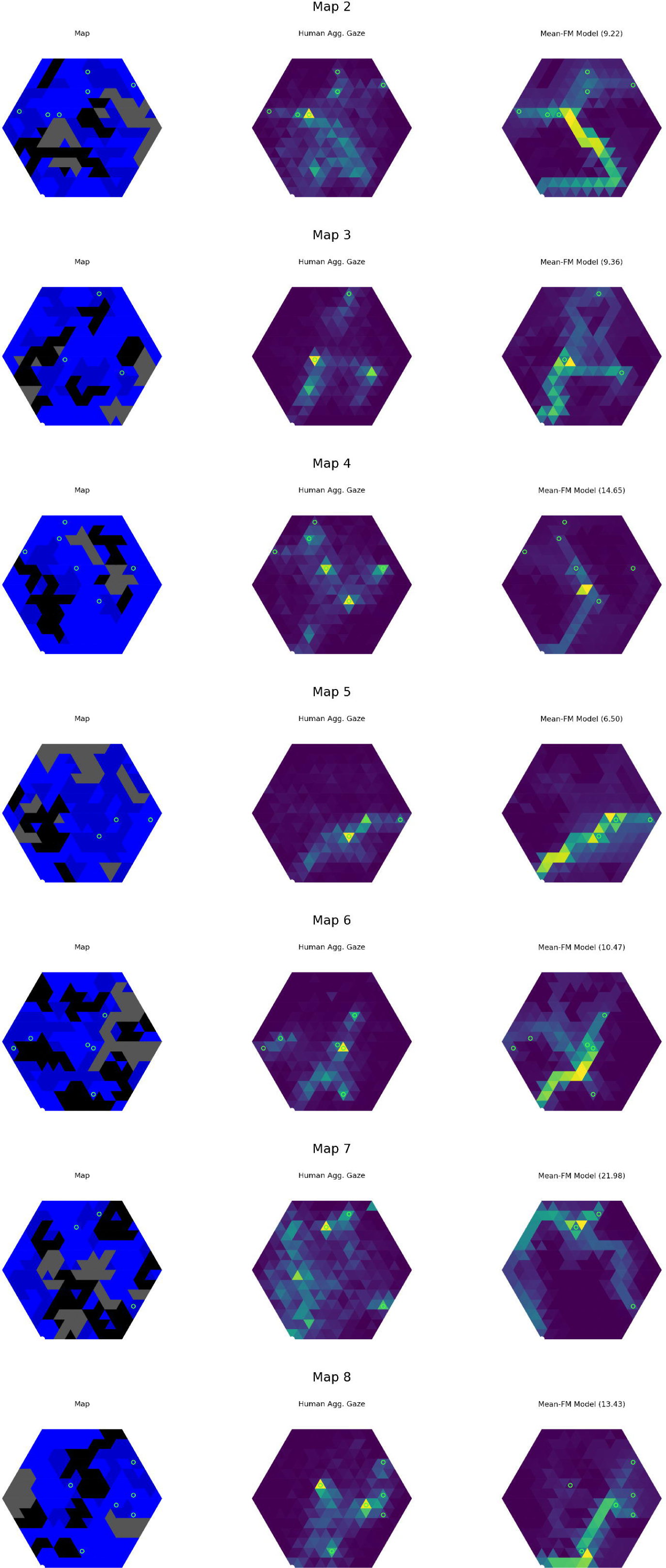
Map-level gaze summary of: a) map geometry, b) aggregate human gaze, c) uniform linear combination of feature maps. Number in title of FM model and agent gaze reports EMD from human aggregate gaze heatmap. Maps 2-8.

**S3 Fig.**
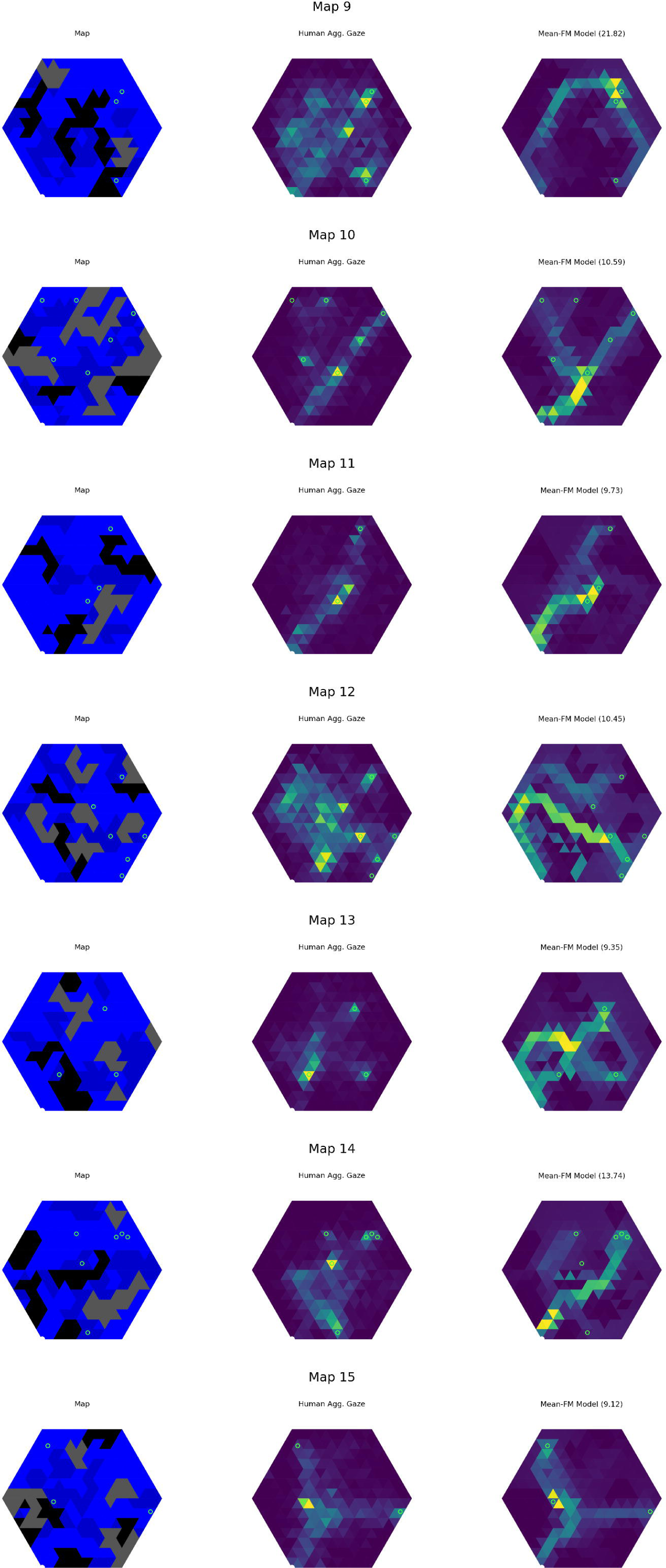
Continuation of S2 Fig. Maps 9-15.

**S4 Fig.**
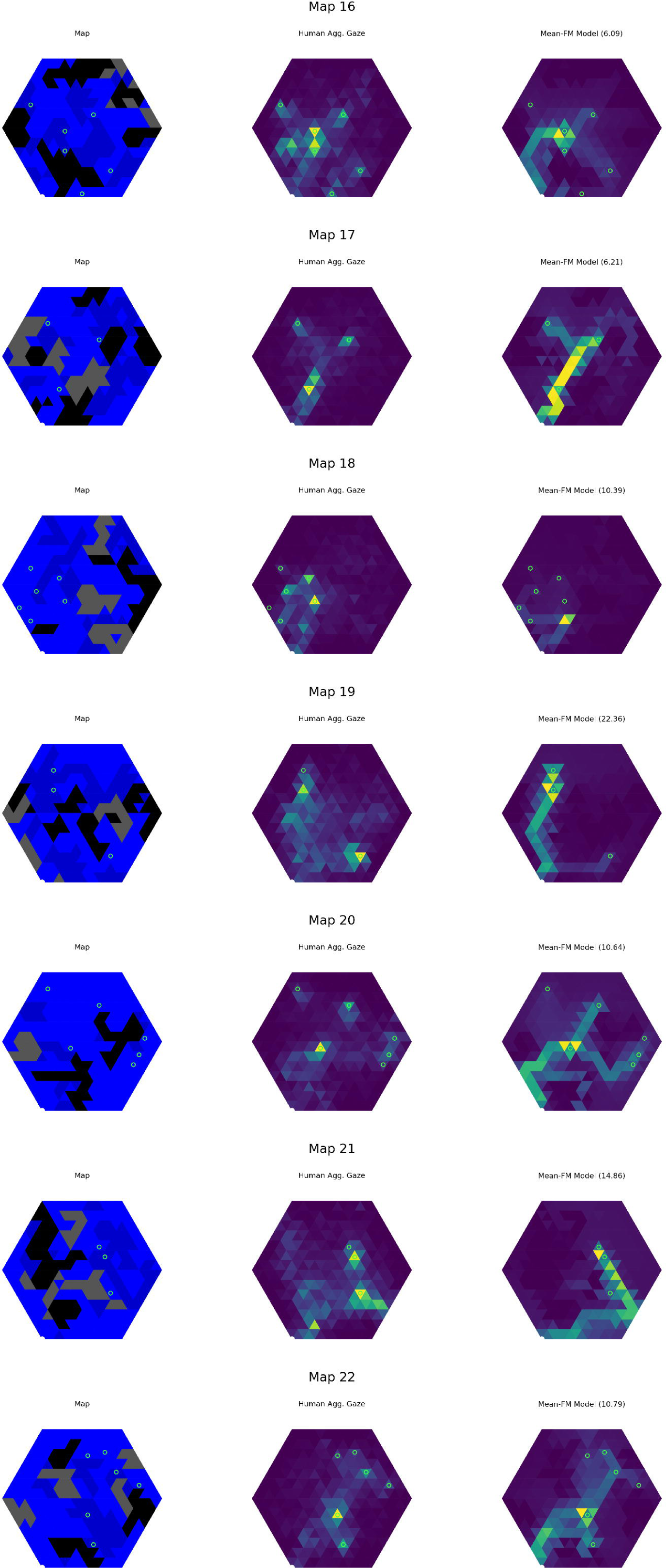
Continuation of S2 Fig. Maps 16-22.

**S5 Fig.**
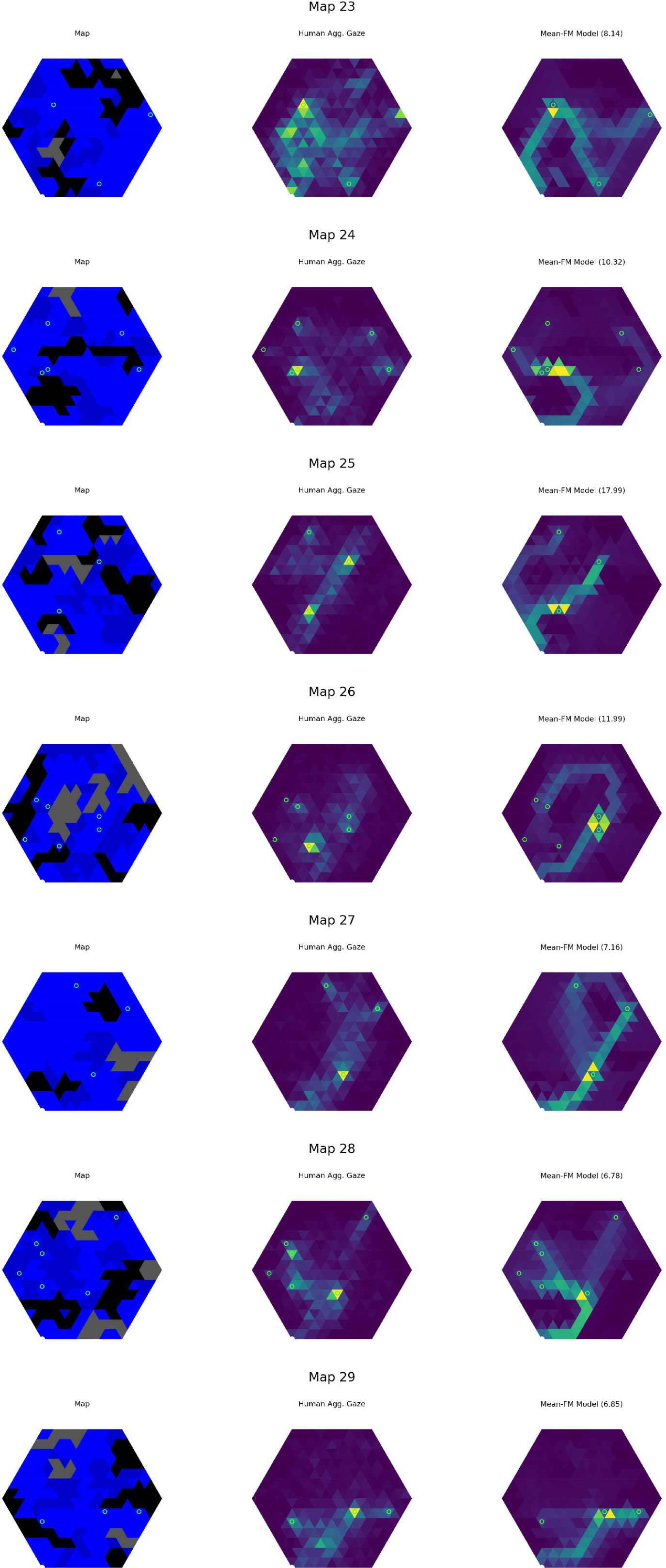
Continuation of S2 Fig. Maps 23-29.

**S6 Fig.**
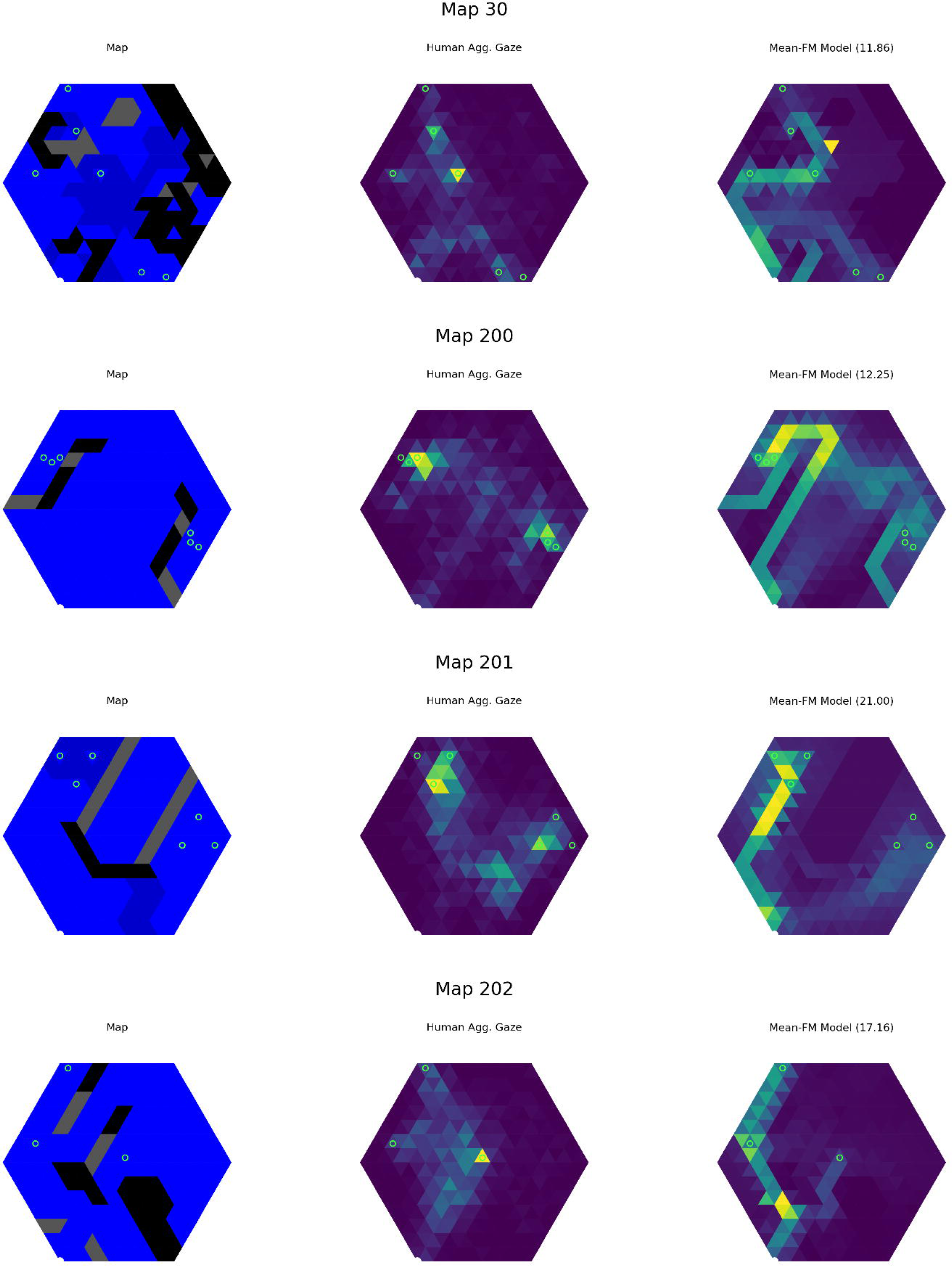
Continuation of S2 Fig. Map 30 and hand designed maps 200-202.

**S7 Fig.**
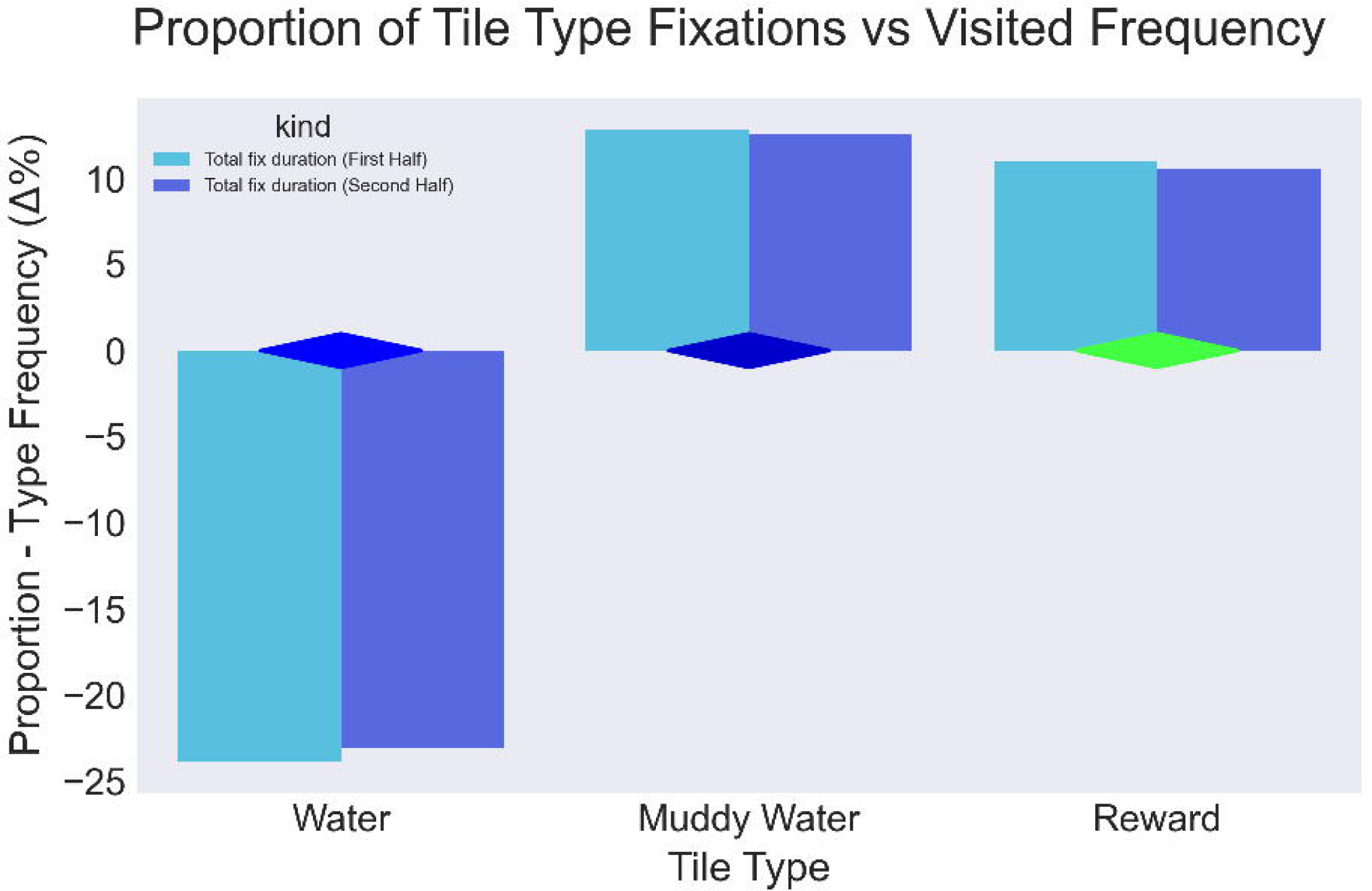
Tile type fixation proportionality vs visitation frequency. Similar analysis to 5 where type frequency is computed based on tile types visited, across participants, during navigation.

**S1 Video. In-task Participant POV.** Short VR video showing sample participant’s perspective during a trial (planning, transition, and navigation phases).

**S2 Video. Aggregate gaze for Map 18.** Animation showing aggregate gaze and navigation trajectories segmented by top and bottom performers for Map 18.

**S3 Video. Aggregate gaze for Map 19.** Animation showing aggregate gaze and navigation trajectories segmented by top and bottom performers for Map 19.

**S4 Video. Aggregate gaze for Map 20.** Animation showing aggregate gaze and navigation trajectories segmented by top and bottom performers for Map 20.

## S1 AAppendix.

### Map generation

Maps for the task were generated in a three step procedure: generate map pool, filter based on criteria, and finally, select an experimental sample. Below, we detail the procedure for each.

### Procedurally generate map pool

Map geometry was defined similarly to [36] but with boundaries defined by tile types, rather than walls drawn along tile edges. Specifically, a hexagonal grid composed of equilateral triangular tiles, and a side length of 7, formed the base map. Each tile could be set to one of four types: water (blue), muddy water (dark blue), land (gray), or obstacle (black). Land tiles were low altitude permitting vision, but not navigational access, while obstacle tiles were tall, and blocked visual access. To generate unique maps with variation in structure and connectivity, the algorithm below was followed:

#### Algorithm 1

Procedural map generation

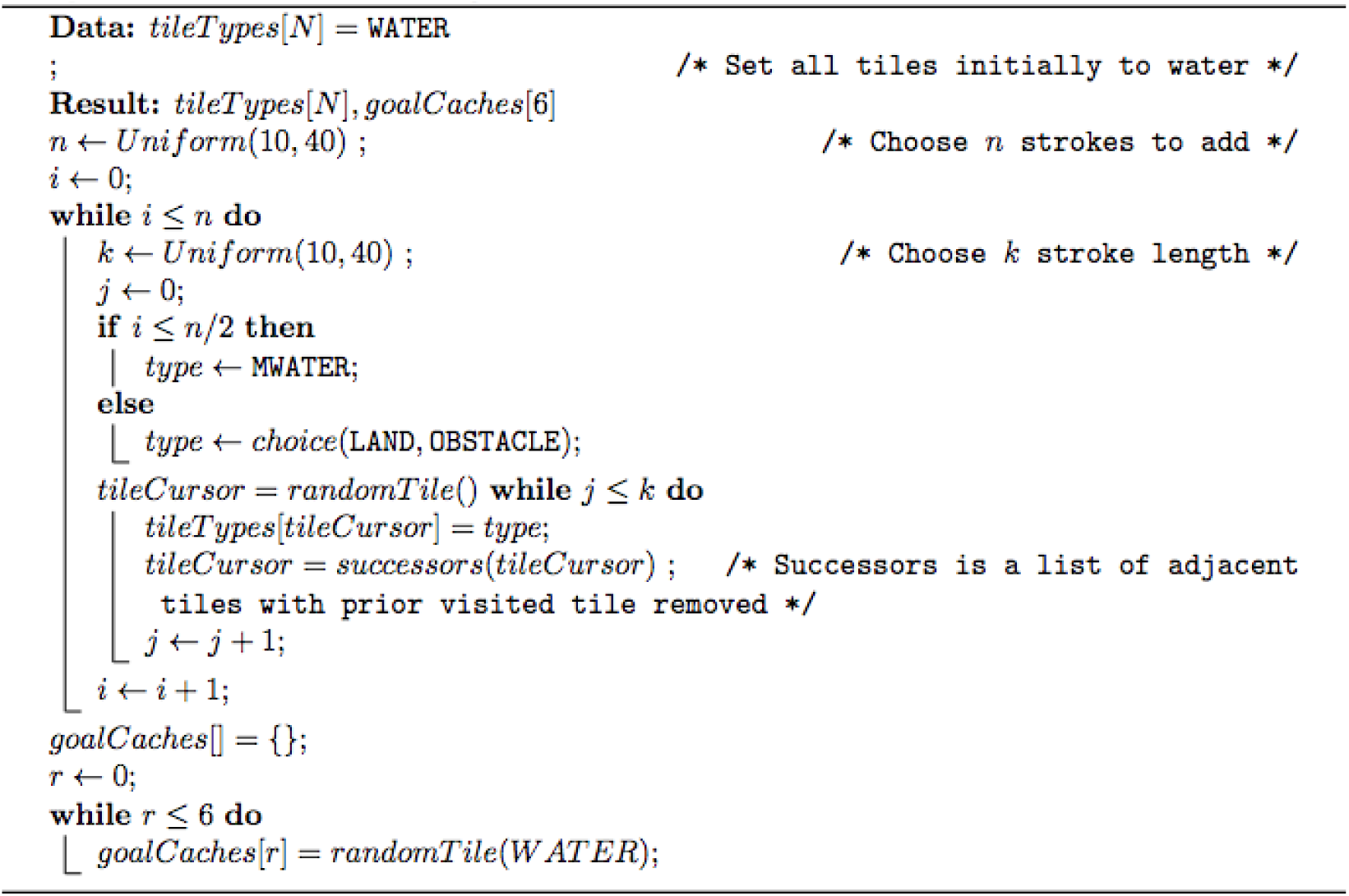

### Filter pool

To filter the pool of map candidates (defined by an assignment for each tile type, and an array of 6 goal cache locations), we confirmed that each one satisfies the following criteria:

1. No goal locations visible (line of sight) from origin
2. All goal locations reachable from origin
3. No obstacles adjacent to origin

### Select experimental sample

Finally, with the filtered pool of eligible map candidates, we selected 30 maps to compose the main experiment. We then randomly assigned half of the maps to each uncertainty group (low vs high). For both groups, we randomly chose 2 goals among the 6 goal caches to be present in the map. For the low uncertainty group, we then randomly removed 3 of the unused goal caches. The end result was a list of maps, where each contained 2 present goals, among either 3 or 6 possible goal cache locations.

Finally, we hand designed two practice maps with trivial geometric structures, to be used at the start of the experiment during the tutorial, as well as three maps testing other information geometries of interest. Map order was randomized for all participants, while ensuring that these latter three maps were always shown at the end of the experiment, to avoid any influence on the primary, procedurally generated maps.

Four sample maps from the main experiment can be seen in S1 Fig (lower-right figures show actual map as represented to participants during planning, with navigation trajectories overlayed).

## S2 Appendix.

While the aim of this study was the way gaze dynamics inform the exploration of many possible plans prior to execution, in light of Ho et. al.’s findings suggesting that individuals pay little attention to task-irrelevant parts of the environment [15], we performed an additional analysis of tile-type fixation proportion that focuses on each individual’s selected trajectory. The analysis was performed in parallel to the tile type analysis shown in Fig 5, however, here we compared fixation duration proportions with visit frequencies across all individuals’ navigated routes (as opposed to the type distribution of the entire map). Results show a similar pattern of relative tile proportionality—less fixation on water, and more on deep water and reward tiles—which helps to validate that the source of the previously reported gaze distribution bias goes beyond overall map structure (see S7 Fig). We note that by restricting the comparison to tiles visited during navigation, the analysis necessarily excludes obstacles and land, which could not be navigated.

## Acknowledgments

This research received funding from the European Research Council under the Grant Agreement No. 820213 (ThinkAhead), the Italian National Recovery and Resilience Plan (NRRP), M4C2, funded by the European Union – NextGenerationEU (Project IR0000011, CUP B51E22000150006, “EBRAINS-Italy”; Project PE0000013, “FAIR”; Project PE0000006, “MNESYS”), and the Ministry of University and Research, PRIN PNRR P20224FESY and PRIN 20229Z7M8N. Support was also received from the UC Berkeley Center for Long-Term Cybersecurity (JG) and the National Science Foundation’s Graduate Research Fellowship Program (JG, Grant No. 2019236659).

## Footnotes

1 The center-line relative position was chosen to capture the ultimate decision of participants to begin navigating to the left or right.

